# A Bidomain Boundary Element–Cable Method for Modeling Neuronal Responses to Electric Fields

**DOI:** 10.64898/2026.09.09.750393

**Authors:** Vanine Sabino, Amanda J. Walenciak, Luis J. Gomez

**Author notes:** Author to whom any correspondence should be addressed. **E-mail:**.

## Abstract

**Objective:** Extracellular electric fields critically influence neural activity through both exogenous neuromodulation and endogenous ephaptic coupling. While conventional cable models efficiently simulate membrane dynamics, they fail to capture bidirectional, field-mediated interactions self-consistently, and fully coupled volumetric methods require computationally prohibitive 3D meshing. We present Cable-BEM, a hybrid wire-kernel bidomain boundary element method designed to resolve these limitations.

**Approach:** By analytically integrating boundary integral kernels around cylindrical neuronal compartments, Cable-BEM fully couples intracellular, extracellular, and membrane dynamics while strictly retaining the highly efficient 1D degrees of freedom of traditional cable equations. The system is advanced using a semi-implicit Crank–Nicolson scheme. To overcome the dense nature of the resulting integral operators, we implement an Adaptive Cross Approximation (ACA) and Hierarchical Off-Diagonal Low-Rank (HODLR) compression scheme.

**Main result:** The solver was rigorously validated against full-surface bidomain boundary element method (BEM) reference implementations, demonstrating tight agreement in activation thresholds (within 1.3% relative error) across diverse stimulation geometries.

The ACA-HODLR compression scheme achieved substantial memory footprint reductions—by a factor of up to 4.6 for large 225-cell networks—without sacrificing numerical accuracy. Furthermore, we utilized the framework to resolve subtle, distance-dependent ephaptic interactions, successfully demonstrating the progressive phase synchronization of biophysically realistic, multi-compartment Purkinje cells.

**Significance:** Cable-BEM provides a computationally scalable, mesh-free framework that establishes a powerful and practical foundation for investigating complex field-mediated phenomena in large-scale, multicellular neuronal networks.

## 1 Introduction

Extracellular electric fields play a fundamental role in both neural stimulation and inter-neuronal communication. In non-invasive and invasive neuromodulation techniques—such as transcranial magnetic stimulation (TMS), transcranial electrical stimulation (TES), and deep brain stimulation (DBS)—externally applied fields are utilized to modulate neural activity for research and therapeutic applications [1, 2, 3, 4]. Conversely, neurons generate endogenous extracellular potentials via their own transmembrane currents, which can subsequently influence adjacent cells through ephaptic coupling [5, 6, 7, 8, 9]. Accurately modeling these reciprocal phenomena requires a computational framework capable of representing membrane dynamics, intracellular conduction, extracellular conduction, and field-mediated cell–cell interactions in a fully self-consistent manner. In this work, we introduce a wire-kernel bidomain boundary element method (BEM) that retains highly efficient, cable-compatible degrees of freedom while seamlessly incorporating fully coupled extracellular, intracellular, and membrane dynamics. By doing so, the resulting formulation upgrades the sparse, nearest-neighbor coupling of the conventional cable equation to encompass nonlocal, field-mediated interactions between all neuronal compartments. To mitigate the computational complexity of the resulting dense operators, hierarchical low-rank representations are employed.

The cable equation remains the most widely adopted approach for simulating neuronal membrane dynamics. By representing neuronal morphologies as interconnected one-dimensional compartments, cable-based methods efficiently simulate action-potential initiation, propagation, and complex conductance-based membrane behaviors [10, 11]. For many stimulation paradigms—particularly when the dominant electric field component is strictly longitudinal to the neurite axis—cable models yield accurate and computationally tractable predictions [12, 13]. However, conventional cable formulations typically treat the extracellular potential as a rigidly prescribed boundary condition or compute it entirely independently of the membrane dynamics. In such hybrid or post hoc workflows, the externally applied field is calculated first and subsequently injected into the cable equation as an extracellular voltage, an activating function, or a distributed source term [13]. Similarly, the extracellular potentials generated by neural activity are frequently approximated post hoc using point-source or line-source models [14, 15, 16]. While historically useful, these unidirectional approaches fundamentally fail to capture the bidirectional feedback loop between the extracellular potential, the transmembrane current, and the intracellular potential.

This limitation becomes critical in scenarios where the extracellular potential serves not merely as an output observable, but as a driving mechanistic variable. Notable examples include ephaptic coupling in densely packed tissues, stimulation within restricted extracellular spaces, transverse polarization effects, and instances where the presence of surrounding cells significantly distorts the device-induced primary field [17, 18]. In standard network simulations, neuronal interactions are predominantly restricted to chemical synapses, gap junctions, or static applied fields. Ephaptic coupling operates via a distinct mechanism: a neuron’s active membrane currents dynamically alter the local extracellular potential, thereby perturbing the membrane voltage of adjacent cells without any direct synaptic or gap-junctional linkage [19]. Recent fully coupled Extracellular–Membrane–Intracellular (EMI) studies indicate that under typical physiological conditions, these ephaptic effects primarily manifest as subtle modulations of spike timing, phase synchronization, and excitability thresholds, rather than as direct triggers for action potentials [20]. Consequently, scalable computational tools are essential to investigate these subtle, yet potentially highly impactful, field-mediated effects across realistic morphologies and large multicell geometries.

Bidomain and EMI-type finite element methods (FEM) provide a rigorous, physically principled approach to modeling these interactions. In such formulations, the intracellular and extracellular spaces are discretized explicitly, and the membrane is modeled as an active interface where voltage-dependent ionic currents continuously couple the two conductive domains [18, 21, 22, 23, 24]. This facilitates a perfectly self-consistent solution for the membrane voltage, transmembrane current, and extracellular potential. However, FEM inherently requires volumetric meshes that must conformably resolve the intracellular space, the bulk extracellular medium, the complex folded membrane surfaces, and the ultra-narrow extracellular clefts separating adjacent cellular structures. For anatomically realistic neuronal morphologies or dense multicell ensembles, this volumetric meshing requirement rapidly becomes a prohibitive computational bottleneck, particularly when micrometer-scale cellular details must be embedded within macroscopic tissue or device domains [25, 26].

Boundary element methods offer an elegant alternative by rigorously reducing the volumetric conduction problem to equivalent unknowns defined exclusively on domain interfaces [27, 28, 29]. Our previous full-surface bidomain BEM formulation demonstrated that membrane dynamics and extracellular conduction can be precisely coupled without necessitating any volumetric meshing [30]. This boundary-centric approach is exceptionally well-suited for stimulation scenarios, as macroscopic tissue boundaries, microscopic cell membranes, and device-induced field sources can be unified within a single boundary integral framework. Nevertheless, directly applying a surface BEM discretization to realistic neuronal morphologies still imposes a substantially higher number of unknowns relative to a 1D cable model, as the entire cylindrical surface area of the complex dendritic arbor must be explicitly tessellated. Furthermore, standard BEM matrices are fully dense, rendering memory storage and repeated temporal stepping computationally exorbitant without the integration of fast algebraic algorithms [31].

To bridge this gap, we introduce a wire-kernel bidomain BEM that strictly preserves the highly efficient compartmental structure of standard cable models while fully retaining the rigorous field coupling intrinsic to a bidomain boundary integral formulation. The central geometric approximation is the assumption that the transmembrane voltage and current remain constant in the azimuthal direction around each localized cylindrical compartment. Rather than explicitly triangulating the complete cylindrical membrane surface, we analytically integrate the boundary integral kernels around the cylinder circumference. This analytical reduction yields a strictly one-dimensional centerline discretization, where the spatial interaction kernels are efficiently evaluated using elliptic integrals. Consequently, each discrete cable compartment contributes the exact same number and type of degrees of freedom as in a standard cable model, yet the rigorous extracellular and intracellular field interactions spanning all network compartments are comprehensively preserved through the governing bidomain integral equation.

The proposed formulation can therefore be understood as a fully cable-compatible bidomain solver. It maintains a computational architecture that is directly analogous to the compartment models universally employed in computational neuroscience, while elevating the local, purely axial coupling of the standard cable equation into a generalized, nonlocal, field-coupled operator. This mathematical framework empowers the method to simultaneously capture macroscopic device-induced fields, microscopic cell-generated extracellular potentials, and dynamic ephaptic interactions within a single, unified time-domain simulation. Because the resulting integral operator is dense, we exploit hierarchical off-diagonal low-rank (HODLR) representations to aggressively compress storage requirements and accelerate computational throughput [32, 33]. Specifically, adaptive cross approximation (ACA) is utilized to construct the low-rank off-diagonal blocks algebraically, completely circumventing the need to assemble the dense interaction matrices explicitly [34, 35, 36, 37]. Such ACA-based hierarchical compression techniques have been extensively validated in large-scale electromagnetic integral-equation systems [37, 38, 39] and have recently been successfully adapted for full-surface bidomain-BEM models of neuronal tissue [31].

We validate the proposed method using canonical neuronal stimulation and action-potential propagation benchmarks, systematically comparing the wire-kernel bidomain BEM against both conventional cable-equation solvers and full-surface bidomain BEM results. These quantitative comparisons verify that the wire-kernel bidomain BEM exactly recovers standard cable theory in regimes where purely axial approximations are valid, while simultaneously unlocking the capability to model complex field-mediated interactions that are fundamentally absent from standard cable formulations. We subsequently deploy this framework to investigate ephaptic coupling and stimulation-induced membrane polarization in multi-neuron configurations. Ultimately, the resulting solver establishes a practical, highly scalable pathway for neuro-simulations that successfully fuses the computational efficiency and familiar morphological degrees of freedom of traditional cable models with the rigorous, self-consistent physical coupling of advanced bidomain and EMI formulations.

## 2 Methods

### 2.1 Cable-Equation Formulation

We first introduce a standard cable-equation formulation, since the proposed boundary-element method is built to use the same compartmental degrees of freedom. In cable solvers, a neuron is represented using a multi-compartment formalism in which the morphology is partitioned into contiguous cylindrical compartments [40, 41, 42, 43]. Each compartment is assigned a length, radius, intracellular resistivity, membrane capacitance, and membrane mechanisms. The membrane mechanisms may include passive leakage, voltage-dependent ion channels, synapses, gap junctions, or prescribed current injections [11, 44]. Branching morphologies are stored either through parent– child relationships or equivalently as a graph of neighboring compartments, as shown schematically in Fig. 1.

**Figure 1.**
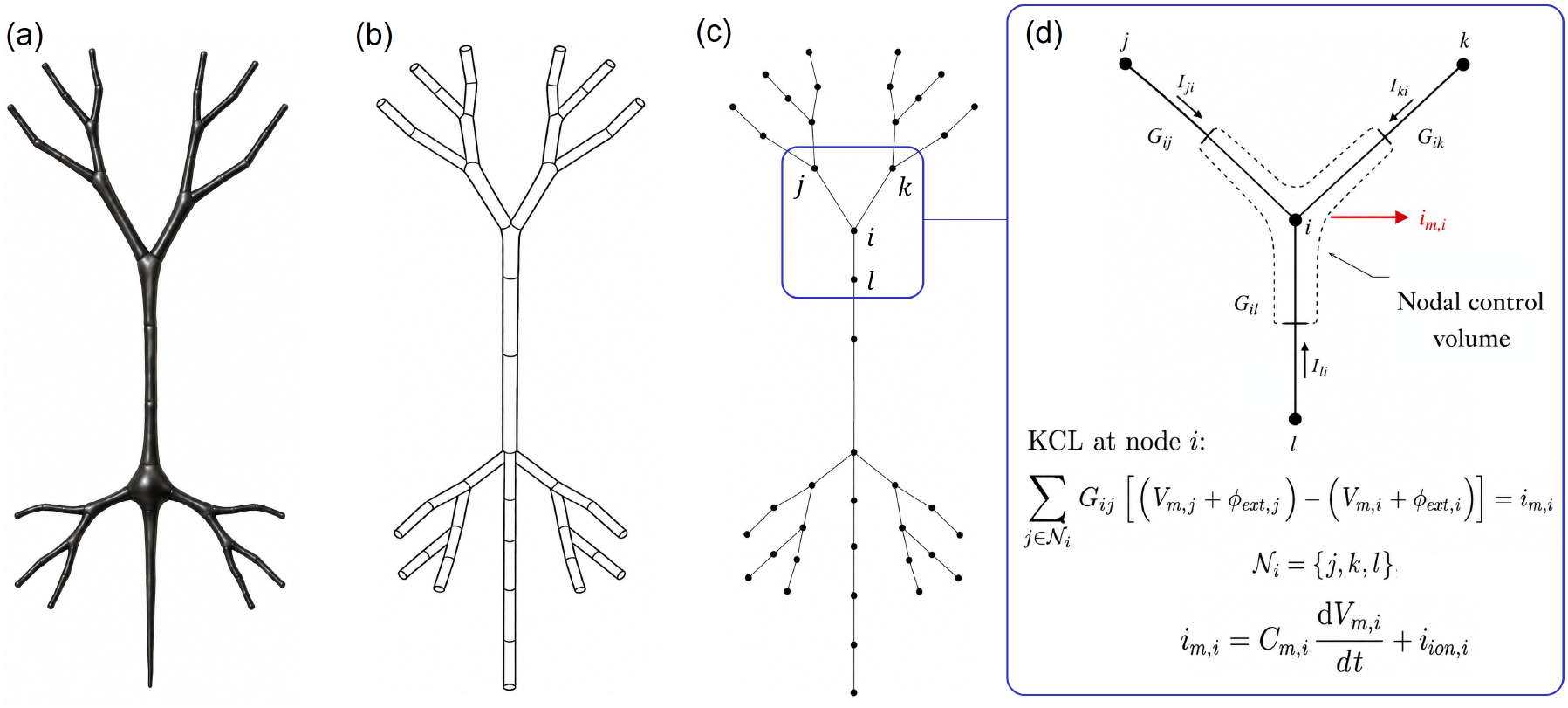
Compartmental cable-model representation and nodal formulation. (a) Example neuronal morphology. (b) Multi-compartment approximation using connected tapered cylindrical segments. (c) Corresponding one-dimensional nodal graph, with nodes located at compartment endpoints and edges representing axial intracellular connections. The highlighted region indicates branching node *i* used to illustrate the local cable formulation. (d) Nodal control volume associated with branching node *i*, bounded by the midpoints of the incident compartments. The connected branches are characterized by the axial conductances *G*_*ij*_ , *G*_*ik*_, and *G*_*il*_, and the corresponding axial currents balance the membrane current *i*_*m,i*_ according to Kirchhoff’s current law. Here, *N*_*i*_ = *{j, k, l}* denotes the set of nodes neighboring *i*.

Schematically, the multi-neuron morphology is represented as a tree graph, where each of the *N*_*s*_ compartments is a vertex in the set *V* , and each edge in the set ℰis a linearly tapered cylinder connecting a compartment to its parent. The cylinder radius varies linearly between the radii of the adjacent compartments. To accommodate changes in compartment properties, each edge stores two compartment-type labels, one associated with each half of the cylinder adjacent to the respective compartment center. This representation permits discontinuities in electrical properties at the mid-point of an edge while maintaining a simple graph structure.

The unknowns are the nodal transmembrane voltages 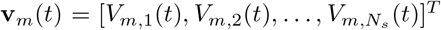 , where *V*_*m*_(**r**, *t*) = *ϕ*_in_(**r**, *t*) − *ϕ*_ext_(**r**, *t*), and *ϕ*_in_ and *ϕ*_ext_ denote the intra- and extracellular potentials, respectively. Conventional cable solvers do not solve for the extracellular potential generated by membrane currents. Instead, *ϕ*_ext_(**r**, *t*) is typically set to ground or prescribed from a separate field calculation when an externally applied field is present [13, 12, 45]. The intracellular potential at each node is therefore written as *ϕ*_in,*i*_(*t*) = *V*_*m,i*_(*t*) + *ϕ*_ext,*i*_(*t*).

The cable approximation assumes that axial current flows only along the cell centerline and that the intracellular potential varies linearly between neighboring compartments. Thus, each graph edge connecting neighboring nodes *i* and *j* is replaced by an axial resistor. For each cylindrical connection, the resistance is 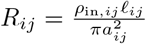 , where *l*_*ij*_ is the length of the cylinder, *a*_*ij*_ is its average radius, and *ρ*_in,*ij*_ is the intracellular resistivity. At each node, Kirchhoff’s current law is applied to a control volume formed by the nearest half of each cylinder incident on that node. In other words, the control volume associated with node *i*, where *i* = 1, 2, … , *N*_*s*_, consists of all half-compartments adjacent to that node. Applying Kirchhoff’s current law to each nodal control volume gives

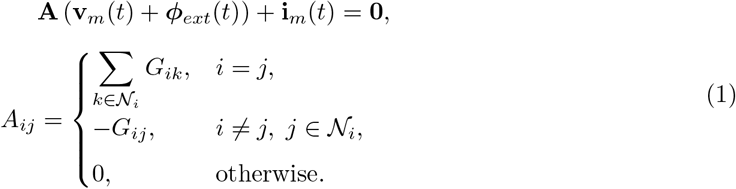

Here, **A** is the weighted graph Laplacian associated with axial intracellular conduction, *N*_*i*_ is the set of compartments neighboring node *i, G*_*ij*_ = 1*/R*_*ij*_ is the axial conductance between neighboring nodes, and **i**_*m*_(*t*), described next, is the net membrane current leaving the control volume. When the extracellular potential is treated as ground, Eq. (1) reduces to **Av**_*m*_(*t*) + **i**_*m*_(*t*) = **0**.

The net membrane current flowing out of node *i* is [**i**_*m*_(*t*)]_*i*_ is obtained by summing the contributions from the portions of all incident edges belonging to that node. If edge *e* represents a cylindrical membrane segment with surface area *S*_*e*_ = 2*πa*_*e*_*l*_*e*_ and approximately constant transmembrane current density *I*_*m,e*_(*t*), then

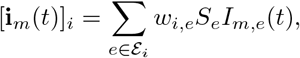

where ℰ_*i*_ is the set of edges incident on node *i*. For the midpoint control volumes used here, *w*_*i,e*_ = 1*/*2 for each endpoint of a cylindrical segment. At branch points, [**i**_*m*_(*t*)]_*i*_ is the sum of the half-segment membrane currents from all connected branches. Lumped regions, such as a soma, are included by adding their assigned membrane area to the corresponding nodal control volume.

The membrane current density consists of two components written as

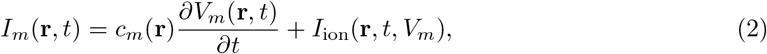

where *c*_*m*_(**r**) is the specific membrane capacitance per unit area and *I*_ion_(**r**, *t, V*_*m*_) is the ionic current density from the assigned membrane model. The ionic current is evaluated from the local membrane mechanisms, typically as a sum of conductance-based currents of the form *g*_*k*_(**r**, *t*)[*V*_*m*_(**r**, *t*) − *E*_*k*_], with the conductances updated through their associated gating variables [10]. The corresponding equivalent circuit at node *i* is shown in Fig. 2, where the membrane current is decomposed into capacitive and ionic components and the neighboring compartments are coupled through axial conductances.

**Figure 2.**
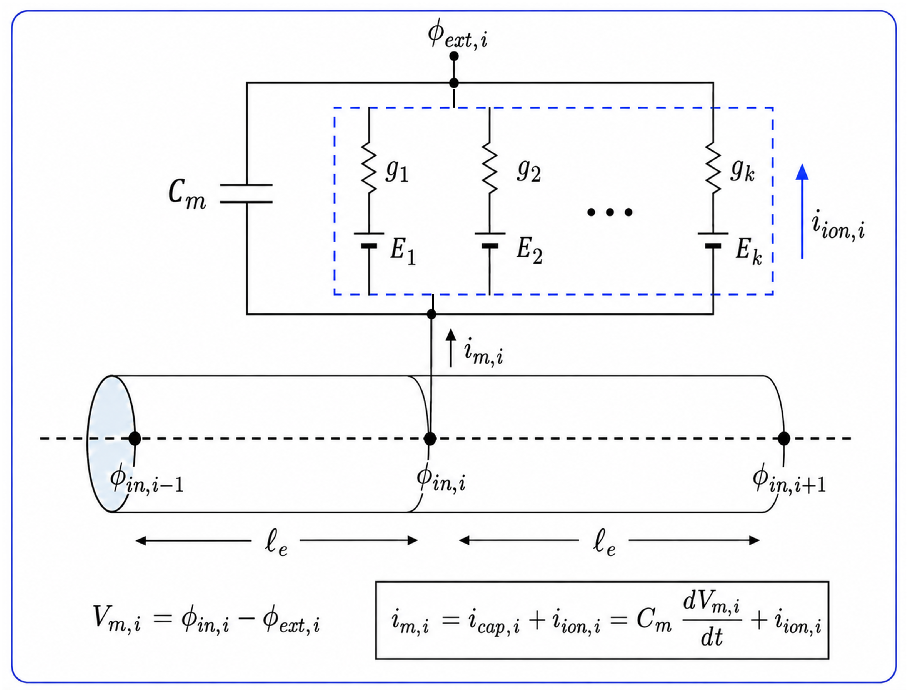
Equivalent circuit associated with node *i* in the cable model. The intracellular potential is represented at nodes located at the endpoints of adjacent cable elements, each of length *l*_*e*_. The membrane current *i*_*m,i*_ is decomposed into a capacitive component, *i*_*cap,i*_ = *C*_*m*_ *dV*_*m,i*_*/dt*, and an ionic component, *i*_*ion,i*_, where the transmembrane voltage is *V*_*m,i*_ = *ϕ*_*in,i*_ *− ϕ*_*ext,i*_. The ionic current is modeled as the sum of parallel conductance-based branches with reversal potentials *E*_1_, *E*_2_, … , *E*_*k*_ and conductances *g*_1_, *g*_2_, … , *g*_*k*_.

Equations (1) and (2) define the cable-equation system used for comparison in this work. In our previous work, we found that cable-equation solutions can agree closely with full bidomain BEM solutions in many cases, suggesting that the main limitation of the cable formulation is not necessarily the compartmental approximation of the neuronal unknowns, but rather the treatment of *ϕ*_*ext*_(*t*) as prescribed instead of solving it self-consistently from the membrane currents [30]. In the following sections, we retain the same compartmental voltage and membrane-current notation and approximations, but replace the prescribed extracellular potential with a bidomain boundary-element formulation that couples the intracellular, membrane, and extracellular fields. This results in a formulation that preserves the simplicity of the cable-equation degrees of freedom while incorporating self-consistent intra- and extracellular field coupling.

### 2.2 Bidomain Boundary-Element Formulation

This section introduces the boundary integral equations used for the bidomain BEM, which will later be discretized using the compartmental approximations introduced above. Only a brief description of the model is given here. Detailed derivations can be found in our previous work [30]. The BEM unknowns are defined on interfaces between regions with distinct electrical properties. We denote the union of these interfaces by Γ, which includes non-membrane conductivity interfaces (Γ_*σ*_), active neuronal membranes (Γ_act_), and electrically insulating membrane regions (Γ_ins_). For each interface, a unit normal vector 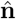 is chosen to point from the− side of the interface to the + side, and the conductivities on the two sides are denoted by *σ*^*−*^ and *σ*^+^. On neuronal membranes, we use the convention *σ*^*−*^ = *σ*_*in*_ and *σ*^+^ = *σ*_*ext*_. This ensures that 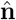points from the intracellular space to the extracellular space, corresponding to the direction of positive transmembrane current.

Under the quasi-static approximation, the electric fields and conduction currents are generated by three contributions: externally applied device fields, charges accumulated on interfaces, and jumps in scalar potential across cell membranes. Electrodes, injected currents, and coils provide prescribed forcing terms. The interface charge density *ρ*(**r**, *t*) generates the secondary electric field. On neuronal membranes, the transmembrane voltage *V*_*m*_(**r**, *t*) = *ϕ*_*in*_(**r**, *t*) − *ϕ*_*ext*_(**r**, *t*) represents a jump in scalar potential and therefore produces an additional field contribution. This membrane voltage term is a zero-thickness representation of the field generated by charge separation across the thin membrane.

The bidomain integral equation is derived by enforcing continuity of normal conduction current across each interface,

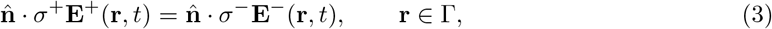

where **E** ^*±*^ (**r**, *t*) denotes the total electric field evaluated on the + and − sides of the interface, respectively. The total field is the sum of the device-induced field, the field generated by interface charges, and the field generated by membrane voltage jumps. Prescribed electrode and coil terms are collected into a known device forcing term.

Substituting the boundary integral representations of these field contributions into Eq. (3) and rearranging gives the bidomain boundary integral equation

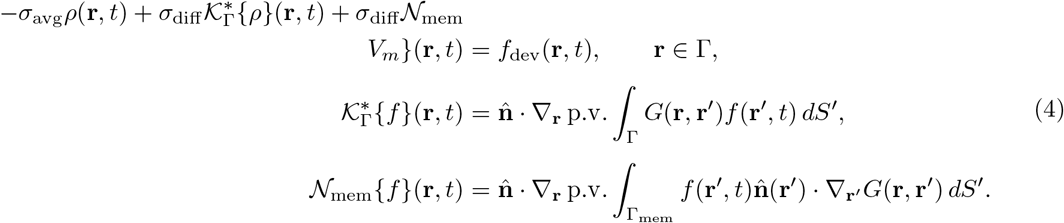

Here, *σ*_avg_ = (*σ*^+^ + *σ*^*−*^)*/*2, *σ*_diff_ = *σ*^+^ − *σ*^*−*^, and *G*(**r, r**′) = 1*/*(4*π*|**r** − **r**′|) is the free-space Green’s function. The term 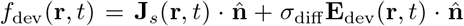 contains the known device-induced currents and primary electric-field contributions, where **E**_dev_(**r**, *t*) denotes the primary device-induced electric field. The operator 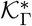 acts on the interface charge density, and *N*_mem_ acts on the voltage jump across neuronal membranes. Thus, the primary unknowns in the bidomain formulation are the interface charge density *ρ*(**r**, *t*) and, on membrane regions, the transmembrane voltage *V*_*m*_(**r**, *t*).

Equation (4) is the continuous boundary equation used throughout the remainder of the formulation. It can be interpreted as Kirchhoff’s current law on the boundary. In other words, the currents generated by device sources, interface charges, and membrane voltage jumps must balance to ensure that the total normal conduction current is continuous across each interface. On neuronal membranes, the interface charge density is proportional to the transmembrane current density. This relation is *I*_*m*_(**r**, *t*) = *η*_*m*_*ρ*(**r**, *t*), where *η*_*m*_ = (1*/σ*_*ext*_ − 1*/σ*_*in*_)^*−*1^. Thus, on active membrane regions, Eq. (2) provides the additional state equation that relates *V*_*m*_(**r**, *t*) to *I*_*m*_(**r**, *t*), and by proxy to *ρ*(**r**, *t*)

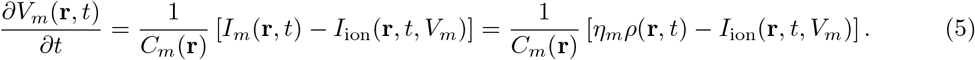

On each interface type, *ρ*(**r**, *t*) and *V*_*m*_(**r**, *t*) satisfy

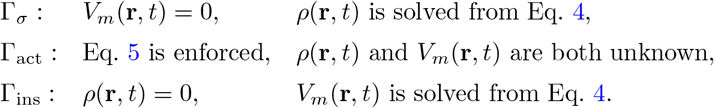

On non-membrane interfaces there is no scalar-potential jump, so *V*_*m*_(**r**, *t*) is set to zero. On insulating membrane regions, no transmembrane current is allowed. Since *I*_*m*_(**r**, *t*) = *η*_*m*_*ρ*(**r**, *t*), this is imposed by setting *ρ*(**r**, *t*) = 0 while still allowing the membrane voltage to polarize through the boundary integral equation.

At junctions between different interface types, *V*_*m*_(**r**, *t*) is required to remain continuous, while *ρ*(**r**, *t*) may be discontinuous. At boundaries between active and insulating membrane regions, the shared value of *V*_*m*_(**r**, *t*) is determined by the neighboring active membrane equation, while the insulating portion enforces *ρ*(**r**, *t*) = 0. At boundaries where a membrane region meets a non-membrane conductivity interface, the non-membrane side enforces *V*_*m*_(**r**, *t*) = 0, correspondingly, the shared membrane voltage unknown is set to zero at that junction. This convention avoids introducing artificial jumps in *V*_*m*_(**r**, *t*) at changes in interface type while preserving the appropriate boundary condition on each region.

### 2.3 Temporal Discretization

Assume that all quantities are known at an initial time *t*_0_. We seek the solution at uniformly spaced time points *t*_*n*_ = *t*_0_ + *n*Δ*t*, for *n* = 1, … , *N*_*t*_, where Δ*t* is the time step. At each step, the device forcing term is evaluated at the new time, 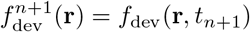.

On active membrane regions, Eq. (5) is advanced using a semi-implicit Crank–Nicolson approximation [18]. The transmembrane current term is approximated using the average of its values at *t*_*n*_ and *t*_*n*+1_, and the ionic current is evaluated using the membrane state from the previous time step:

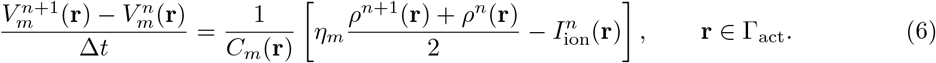

Here, 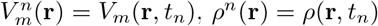, and 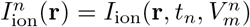. Equivalently,

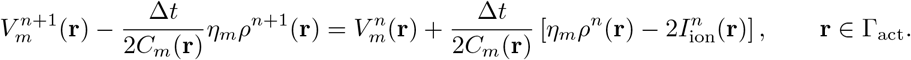

Combining this update with Eq. (4) gives the time-discrete bidomain BEM system of equations.

At each time step, we solve for *ρ*^*n*+1^(**r**) and 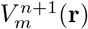 such that

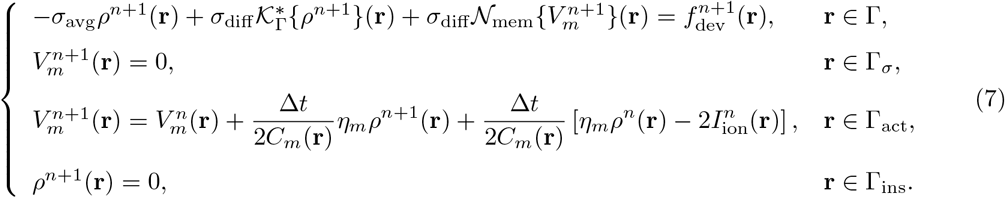

The first line enforces current continuity on all interfaces. The remaining lines impose the appropriate auxiliary condition for each boundary type: no voltage jump on non-membrane conductivity interfaces, the membrane state equation on active membrane regions, and zero transmembrane current on insulating membrane regions.

After solving Eq. (7), the membrane current on active membrane regions is recovered from 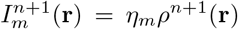, the membrane voltage is updated using Eq. (6), and the ionic state variables are updated using their respective membrane models. In the next section, the cylindrical cable elements are used to discretize *ρ*(**r**, *t*) and *V*_*m*_(**r**, *t*) and obtain the final matrix equations solved at each time step.

### 2.4 Spatial Discretization

We now discretize the time-discrete bidomain equations in space using the tapered cylindrical cable elements illustrated in Fig. 3. The cable equation approximates the membrane voltage as varying linearly along the centerline of each compartment and as constant in the azimuthal direction. Since the active membrane equation linearly relates *V*_*m*_ and *ρ*, we use the same expansions for both quantities.

**Figure 3.**
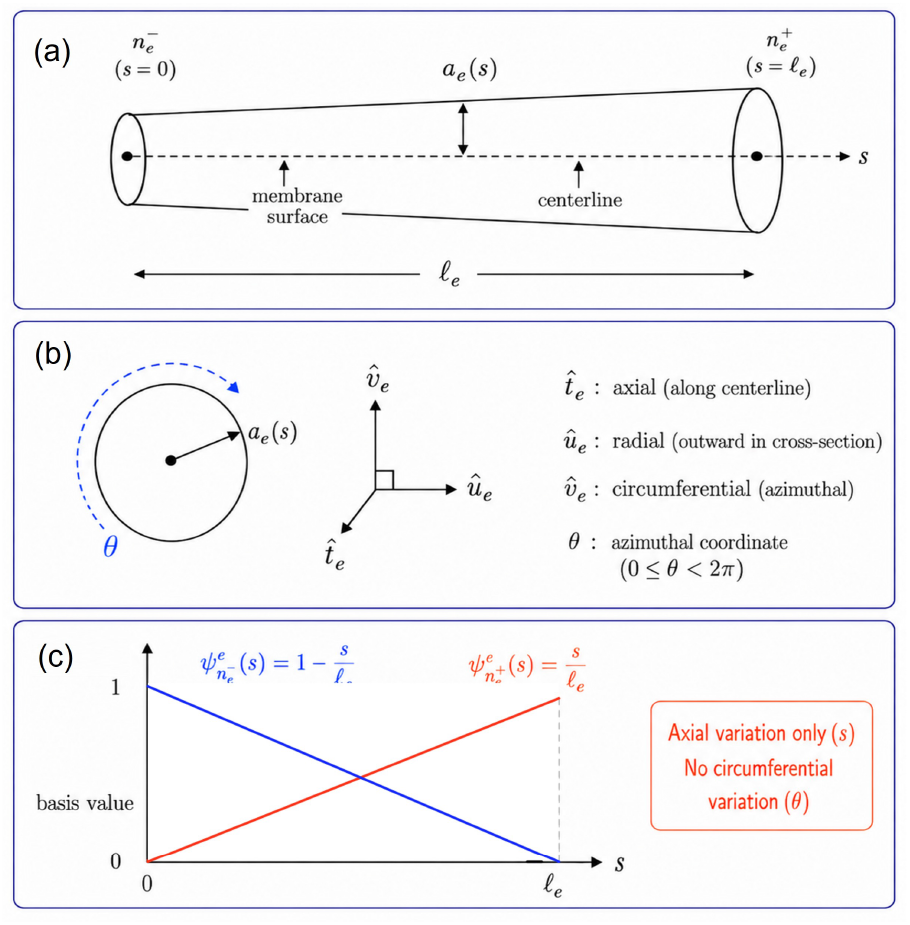
Geometric and basis-function representation of a tapered cylindrical cable element. (a) A cable element of length *l*_*e*_, parameterized by the axial coordinate *s ∈* [0, *l*_*e*_], with endpoint nodes 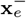and 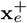 and a linearly varying radius *a*_*e*_(*s*). (b) Local cylindrical coordinate system associated with the element, defined by the axial, radial, and circumferential unit vectors 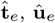, and 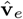, respectively, with azimuthal coordinate *θ*. (c) Linear nodal basis functions 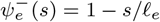and 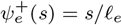 used for interpolation along the axial direction. The approximation varies only along *s* and is assumed uniform in the circumferential direction.

Consider a tapered cable element, or graph edge, *e* connecting adjacent compartment nodes with indices 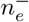 and 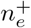. Let 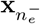and 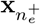 denote the compartment center locations, 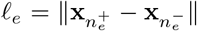 its length, and 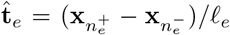 its tangent direction. The circular cross-section radius varies linearly along the cell as 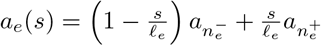 , where 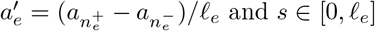 is the coordinate along the centerline.

We choose two orthonormal vectors û_*e*_ and 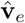spanning the plane normal to 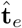_*e*_, with 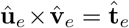. Points on the membrane surface Γ_*e*_ are parameterized as

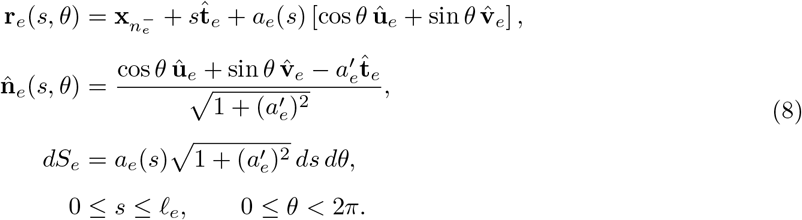

Here, 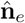 is the outward normal from the intracellular space to the extracellular space. When 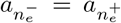, the element reduces to a constant-radius cylinder, 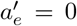, and the normal becomes purely radial. This parameterization makes the azimuthal symmetry of the cable approximation explicit while allowing tapered neuronal sections.

As in the cable equation, we assign one degree of freedom per compartment and expand *ρ*^*n*^(**r**) and 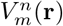 using basis functions 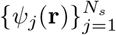 . Each basis function *ψ*_*j*_(**r**) is associated with a single compartment with node index *j* and is nonzero only on the membrane boundary of the cable elements incident on that node. The global nodal basis function *ψ*_*j*_ is assembled by adding all local basis-function contributions that share the same global node index. Thus, ifℰ _*j*_ is the set of cable elements incident on node *j*, then

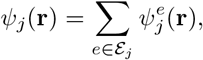

where, on element *e*,

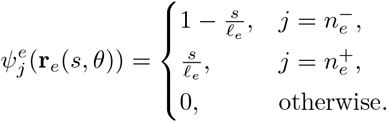

After assembly, *ψ*_*j*_ has the usual hat-function shape along the cable segment and is constant around the circular cross-section. In other words, it is one at node *j*, decreases linearly to zero along each connected cable element, and is zero at the neighboring nodes. To ensure continuity of the approximated quantity, the same nodal coefficient is shared by all incident elements at branch points. Using the above definitions, the cable-consistent approximations are

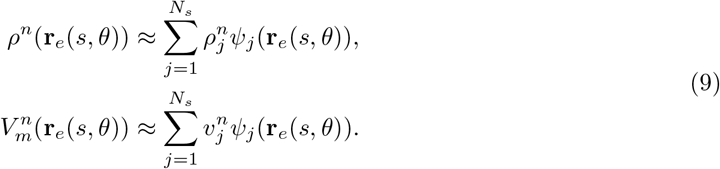

The vectors of coefficients at time *t*_*n*_ are denoted by 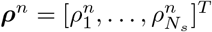and 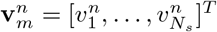With the cable-element parameterization, the source integration over a tapered membrane element separates into an axial integral and an azimuthal integral. For a generic element *e* and local basis function 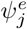, the single- and double-layer potentials generated by that element are

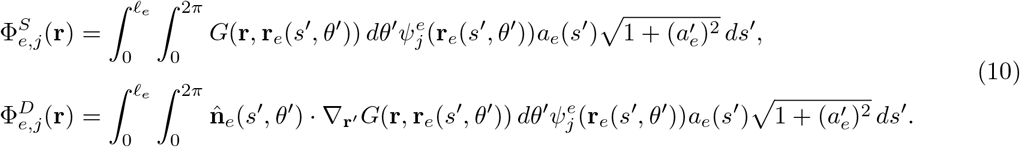

For a global basis function *ψ* _*j*_, the full potential is obtained by summing Eq. (10) over all incident elements. The azimuthal part of the single-layer potential can be evaluated analytically using elliptic integrals [46]. We extended this wire-kernel treatment to the double-layer potential; the formulas used for both kernels are given in Appendix A.

Substituting Eq. (9) into the time-discrete bidomain equation gives the spatially discrete system. The unknowns are grouped according to interface type. Non-membrane interfaces Γ_*σ*_ contribute only charge unknowns, active membranes Γ_act_ contribute charge unknowns, and insulating membranes Γ_ins_ contribute membrane voltage unknowns. Thus, let 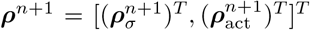and let 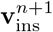 denote the free insulating voltage unknowns.

The active membrane voltage is not kept as an independent unknown. Instead, the membrane state equation is enforced by nodal collocation. At an active membrane node *j*, we use 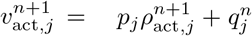, where *p*_*j*_ = Δ*tη*_*m,j*_*/*(2*C*_*m,j*_) and 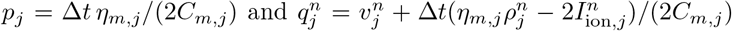. With this substitution, applying the Galerkin procedure to the boundary integral equation gives

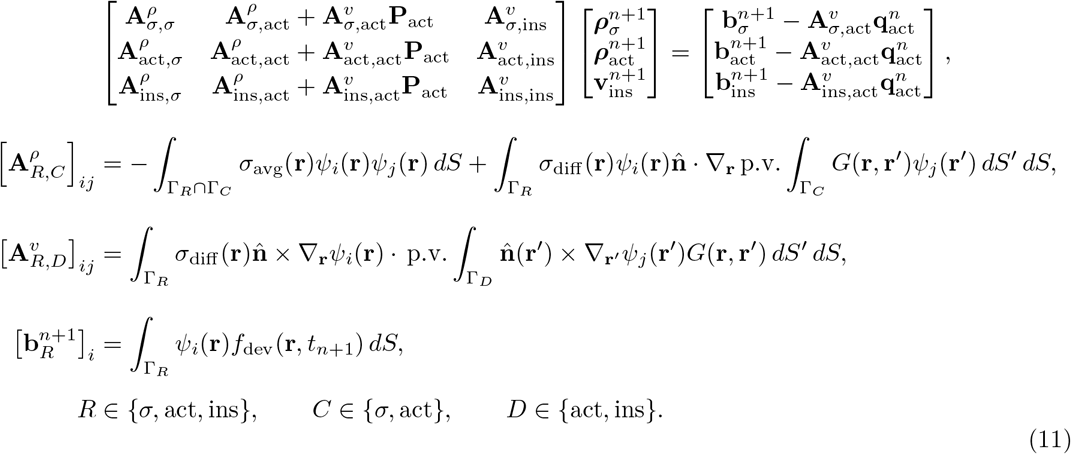

Here **P**_act_ is a diagonal matrix with entries *p*_*j*_ and a vector with entries 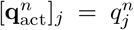. The implementation follows the supports implied by the interface conditions. Charge source integrals are evaluated only over Γ_*σ*_ ∪Γ_act_, since *ρ* = 0 on insulating membrane regions. Membrane voltage source integrals are evaluated only over membrane regions, since *V*_*m*_ = 0 on non-membrane interfaces.

Junction constraints are applied before assembling the matrix by modifying the corresponding basis-function supports and index sets. In particular, at active–insulating membrane junctions, such as transitions to myelinated regions, *V*_*m*_ is kept continuous and the insulating-side value is constrained to the active membrane value determined by the collocated membrane update. Therefore, independent insulating voltage unknowns are introduced only on insulating nodes that are not constrained by neighboring active membrane nodes. In the geometries considered here, membrane interfaces do not share nodes with non-membrane conductivity interfaces; as a result, no additional membrane–non-membrane junction treatment is required. Such treatment would require membrane-side voltages to be constrained to the non-membrane *V*_*m*_ = 0 condition.

After solving Eq. (11), 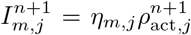 is computed at active membrane nodes, and 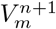 is obtained from the state equation and the insulating membrane solution. Finally, gating variables are updated using the new membrane voltage values.

For each source element, the inner surface integrals in Eq. (11) are evaluated using the tapered cable-element parameterization in Eq. (8). For nearby source and test elements, the azimuthal integrations are evaluated using analytic wire-kernel formulas based on elliptic integrals, provided in Appendix A, and the remaining axial and testing integrations are performed numerically. For well-separated source and test elements, the matrix entries are computed solely using quadrature. To improve the accuracy of the solution, conservation of charge is imposed using an orthogonal projection that ensures charge neutrality at each time step. Finally, the cable BEM does not incorporate transverse stimulation mechanisms, as these would require nonuniform azimuthal variation of the field. These effects are typically negligible and, when present, are incorporated using the approach described in [17].

### 2.5 Fast factorization of system matrix

The Cable-BEM discretization results in a dense system matrix whose storage and factorization costs increase rapidly with the number of unknowns. To reduce these costs, we apply the hierarchical off-diagonal low-rank (HODLR) approach used in [31] to the Cable-BEM system. In the HODLR representation, the matrix is recursively organized into blocks, with the diagonal blocks at the finest level stored explicitly as dense matrices and the off-diagonal blocks describing interactions between sibling groups represented in low-rank form [32, 33]. This hierarchical structure provides a compressed representation of the Cable-BEM system matrix that can subsequently be factorized and used for repeated linear-system solutions. The hierarchical compression has been described in detail in [31]. Next we provide a brief summary with important parameters to this work.

First, each Cable-BEM unknown is associated with a spatial coordinate. The lateral unknowns use the coordinates of the corresponding cable nodes, while the two terminal-cap unknowns of each cell are assigned the coordinates of the corresponding cap centers. The unknowns are then recursively partitioned using a KD-tree. At each level, every group is divided into two child groups by bisecting its axis-aligned bounding box along the coordinate direction with the largest spatial extent. This procedure is repeated for all groups at each level of the hierarchy. The resulting KD-tree defines the hierarchical groups, and the corresponding permutation places the unknowns within each group in contiguous matrix-index ranges [31].

For a Cable-BEM system with *N*_red_ unknowns and a target leaf-size parameter *n*_leaf_ , the number of HODLR levels is chosen as

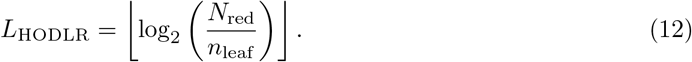

Here, *N*_red_ denotes the total number of unknowns in the Cable-BEM system. The KD-tree partitioning is carried out for *L*_HODLR_ levels, producing 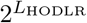leaf groups whose corresponding diagonal matrix blocks are stored densely.

At each level of the HODLR hierarchy, the off-diagonal blocks corresponding to interactions between sibling groups are approximated in low-rank form. For an *m* ×*n* off-diagonal block **A**_*ij*_ ∈ ℝ^*m×n*^, we write

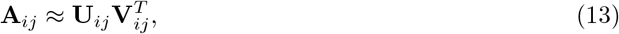

where 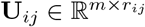 and 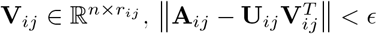, and *ϵ* is chosen to ensure that the end results achieve full accuracy. This results in lower storage whenever *r*_*ij*_ *<* 2 max(*m, n*) as the total storage becomes *r*_*ij*_(*m* + *n*) vs *m* × *n*.

To construct **U**_*ij*_ and 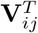 we use a cross-approximation, and a good cross approximation is determined by computing *r*_*ij*_ rows and columns following the adaptive cross-approximation algorithm (ACA) [34, 36, 37]. The ACA approximation rank is estimated from the Frobenius norm of the accumulated low-rank approximation, and the rank is chosen to ensure that the error is below *ℰ*_ACA_. Only the rows and columns requested by the ACA procedure are evaluated; as a result, this also improves the total matrix computation time. Similarly, the cost to factorize and apply the system to a vector is lower using HODLR.

After the HODLR representation is constructed, the system matrix is factorized in HODLR form as

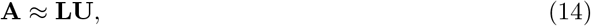

where **L** and **U** are block lower- and upper-triangular HODLR factors, respectively. At the finest level of the hierarchy, each dense diagonal leaf block is factorized using a direct LU decomposition. The factorization then proceeds through the hierarchy by incorporating the low-rank interactions between sibling groups through Schur-complement updates. The full algorithm is described in our previous work [31] and not given here. The resulting low-rank updates are recompressed to maintain the HODLR structure. Once the factors are constructed, repeated Cable-BEM systems with different right-hand sides are solved using forward and backward substitution.

### 2.6 Cell models

We consider several cell types: passive, Hodgkin–Huxley (HH), and Purkinje cell models. For both Passive cell and HH membrane types *c*_*m*_ = 1*µ*F*/*m and the resting voltage is *E*_rest_ = −65 mV. For the passive membrane type *I*_ion_ = *g*_*L*_ (*V*_*m*_ −*E*_*L*_) where *g*_*L*_ = 3S*/*m^2^ is the specific leak conductance and *E*_*L*_ = −65 mV is the leak reversal potential. For Hodgkin–Huxley [10], the ionic current density is 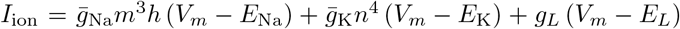, where 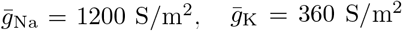, and *g*_*L*_ = 3 S*/*m^2^ are the maximum sodium, potassium, and leak conductances, respectively, and *E*_Na_ = 50 mV, *E*_K_ = −77 mV, and *E*_*L*_ = −54.4 mV are the corresponding reversal potentials. The variables *m, h*, and *n* denote the voltage-dependent gating variables described in [10]. For the Purkinje cell simulations, gates provided by [20] in their supplemental material were compiled and used. Furthermore, the cell morphology and scenario was designed to be similar to the example provided in [20]. Our gate models are all updated using the following formalism

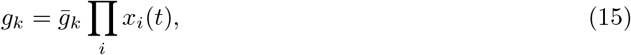

where 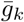 is the maximum conductance of the *k*-th ion channel and *x*_*i*_ is the probability that the *i*-th gating variable is open. The gating variables follow first-order dynamics given by

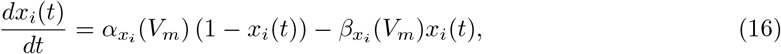

where the rate functions 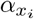 and 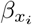 are specific to each channel type.

After the membrane voltage is computed at each time step, the gating variables are updated according to

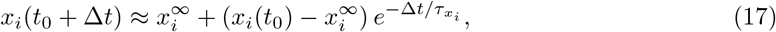

where 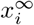 is the steady-state value of the *i*-th gating variable and 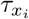 is its time constant.

### 2.7 Simulation Setup and Computational Workflow

Unless stated otherwise, all simulations employ the common electrical and numerical parameters listed in Table 1. The intracellular and extracellular media are assumed to be homogeneous and isotropic. Membrane dynamics are initialized at the resting potential and advanced using the semi-implicit temporal discretization described in Section 2.3. Geometry-specific dimensions, stimulation conditions, intercellular separations, and sampled membrane locations are provided together with the corresponding numerical experiments.

**Table 1.** Common electrical and numerical parameters used in the simulations.

| Parameter | Symbol | Value |
| --- | --- | --- |
| Intracellular conductivity | $\sigma_{\text{in}}$ | 1 S/m |
| Extracellular conductivity | $\sigma_{\text{ext}}$ | 2 S/m |
| Membrane capacitance | $c_m$ | $10^{-2}\text{ F}/\text{m}^2$ |
| Time step | $\Delta t$ | $10^{-5}\text{ s}$ |
| Resting potential | $E_{\text{rest}}$ | $-65\text{ mV}$ |

The computational workflow of the Cable-BEM formulation is summarized in Fig. 4. At the beginning of each time step, the membrane voltage 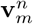, active-membrane charge density 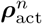, ionic current 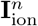, and gating variables **g**^*n*^ are known. These quantities, together with the device forcing at *t*_*n*+1_, are used to assemble the right-hand side **b**^*n*+1^ of the spatially discrete BEM system. The resulting linear system is then solved for the charge density 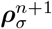 on non-membrane conductivity interfaces, the active-membrane charge density 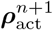, and the insulating-membrane voltage 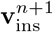. The active-membrane current is recovered as 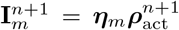, and the active-membrane voltage 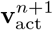 is obtained from the semi-implicit membrane-state relation. The complete membrane-voltage vector 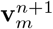 is then assembled from the active- and insulating-membrane values and used to update the gating variables 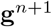 and evaluate the ionic current 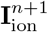. These updated quantities define the state for the next time step.

**Figure 4.**
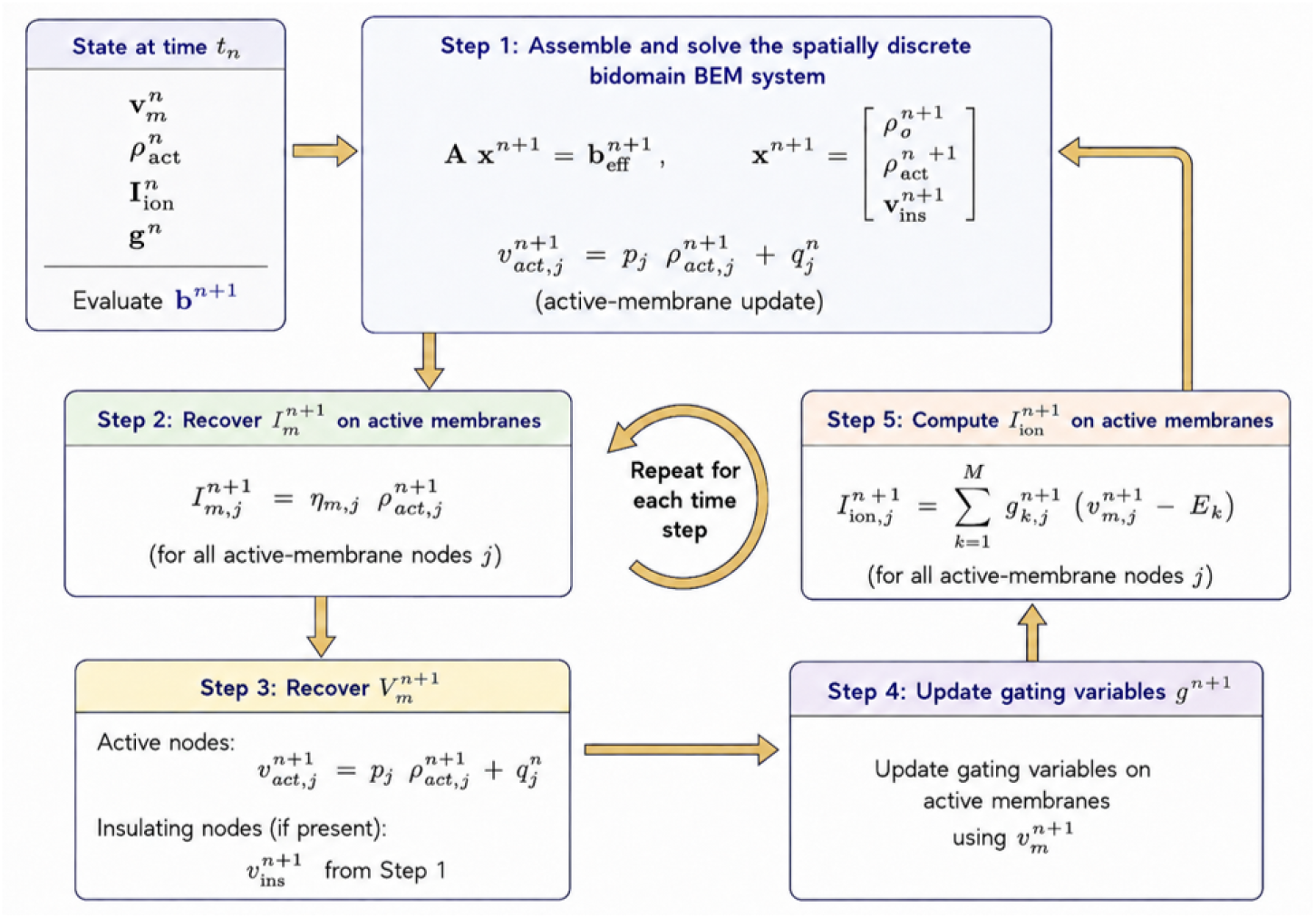
Procedural flowchart of the spatially discrete Cable-BEM algorithm. The sequence is repeated at each time step to update the membrane and ionic state variables.

### 3 Results and Discussion

### 3.1 Validation of the Cable BEM Model

The Cable BEM formulation was validated against established full bidomain-BEM results using three representative benchmarks involving intracellular current injection, stimulation by an externally applied electric field, and multicell field interactions. In each comparison, the membrane properties, stimulation conditions, and temporal discretization were matched between the Cable BEM and the corresponding reference model to ensure that any observed differences were primarily attributable to the spatial formulation.

Figure 5 (a) compares the Cable BEM and Triangle BEM formulations for a straight Hodgkin– Huxley cable subjected to intracellular current injection. The predicted activation thresholds are 71.620 pA and 70.750 pA, respectively, yielding a relative difference of approximately 1.2%. The waveform comparisons shown in the insets further demonstrate close agreement in both the sub-threshold (*I*_inj_ = 45 pA) and suprathreshold (*I*_inj_ = 90 pA) regimes, accurately capturing the propagation delay along the cable.

**Figure 5.**
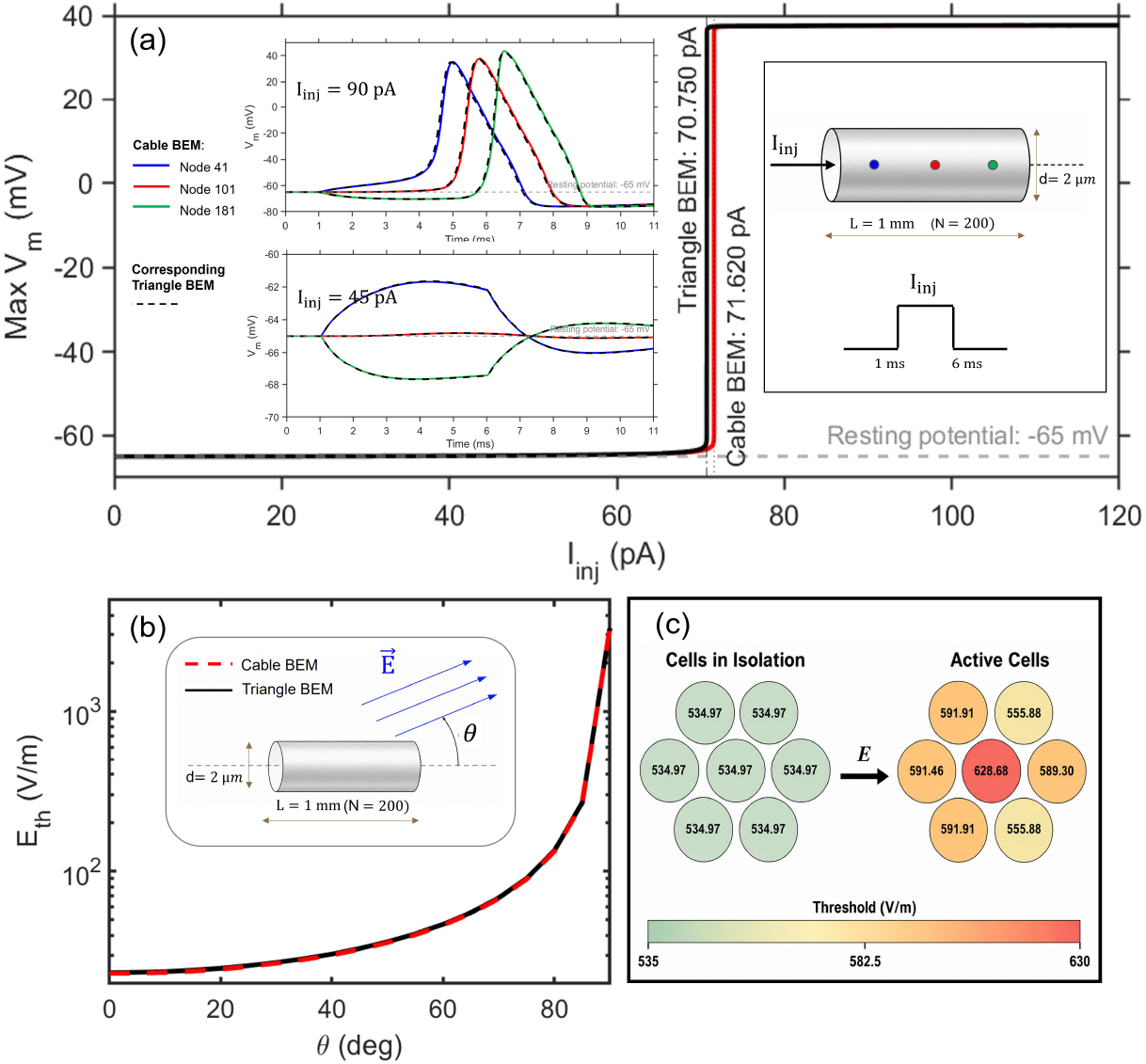
Validation of the Cable BEM formulation against established bidomain-BEM results. **(a)** Current-injection benchmark for a cylindrical Hodgkin–Huxley cable, comparing activation thresholds and representative subthreshold and suprathreshold membrane-potential responses obtained with Cable BEM and Triangle BEM. **(b)** Activation threshold *E*_th_ as a function of the angle *θ* between a spatially uniform applied electric field and the axon axis. **(c)** Activation thresholds for an isolated spherical cell and a seven-cell cluster under a uniform electric field, illustrating the threshold shift produced by neighboring cells.

Figure 5(b) examines stimulation by a spatially uniform electric field. Across field orientations ranging from *θ* = 0° to 85°, the Cable BEM activation threshold differs from the Triangle BEM prediction by less than 1.33%, with a mean relative difference of approximately 1.20%. For the limiting transverse case (*θ* = 90°), the approximation of [17] is employed, and the discrepancy decreases to approximately 0.08%. The threshold for a field perpendicular to the axon is approximately 142 times larger than that for an axially aligned field in the Cable BEM formulation (144 times in the Triangle BEM formulation). This indicates that transverse polarization is only important for edge cases like neurite segments oriented nearly perpendicular to the applied field.

Finally, Figure 5(c) evaluates the formulation using the multicell spherical benchmark from [30]. The isolated-cell activation threshold is *E*_thr_ ≈ 534.97 V*/*m, whereas the threshold of the central cell in the seven-cell cluster increases to 628.68 V*/*m (a 17.5% increase). The corresponding full bidomain-BEM study reported an increase of approximately 15.8%, with both formulations predicting the largest threshold shift for the central cell. Taken together, these benchmarks demonstrate that the reduced Cable BEM formulation successfully reproduces established full-BEM predictions across single-cell stimulation and multicell field-interaction configurations.

### 3.2 Numerical Accuracy and Computational Performance

The numerical behavior of the Cable-BEM formulation was evaluated in terms of spatial convergence, ACA/HODLR compression accuracy, sensitivity to cell packing, and computational scaling with increasing system size. These studies were designed to identify the spatial and compression parameters required for accurate multicellular simulations and to quantify the computational efficiency of the compressed formulation.

#### 3.2.1 Spatial Discretization and Convergence

To assess the spatial resolution required to capture interactions between neighboring cells, convergence studies were performed for two parallel cylindrical cells using active Hodgkin–Huxley and passive membrane models. In each case, the solution obtained with *N* = 1600 compartments per cell was used as the reference, and the relative error was computed as ∥ *x*_*N*_−*x*_1600_ ∥ _2_*/* ∥ *x*_1600_∥ _2_.

Figure 6 presents the active-cell configuration. The error in the directly stimulated cell is several orders of magnitude smaller than the error in the induced response of the neighboring cell across the entire discretization range. All errors decrease monotonically with increasing spatial resolution. For the induced response, the *d* = 0.1 *µ*m and *d* = 1 *µ*m cases exhibit the largest errors, whereas the *d* = 10 *µ*m configuration converges more readily. Thus, resolving the field-mediated response of a neighboring cell is substantially more demanding than resolving the directly stimulated cell, particularly for closely spaced geometries.

**Figure 6.**
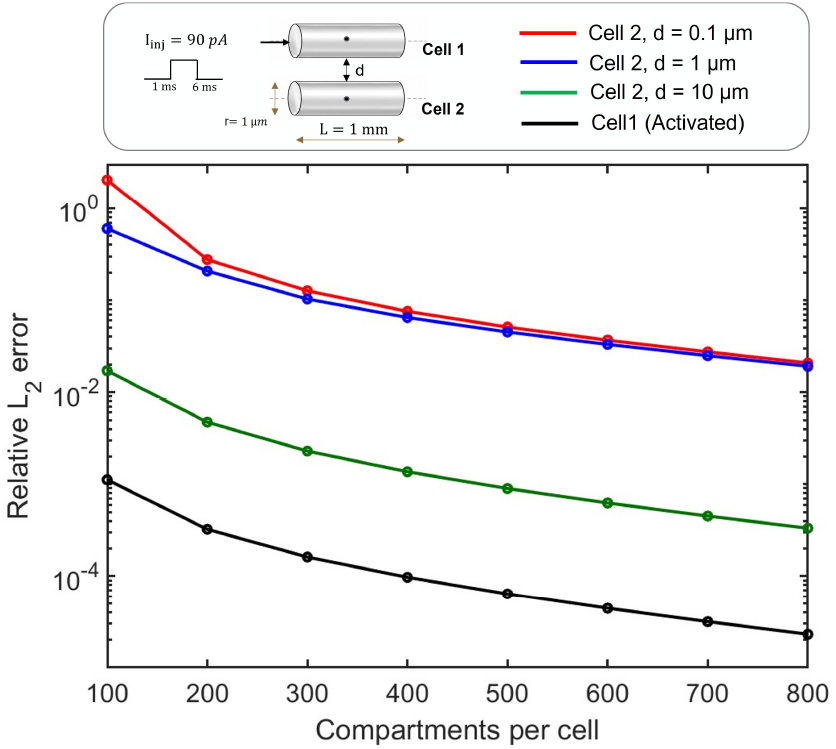
Spatial convergence of the Cable-BEM solution for two parallel active cylinders. Relative *L*_2_ errors were computed with respect to the reference solution obtained using 1600 compartments per cell. The black curve shows the error in the activated cell (Cell 1), while the colored curves show the error in the induced response of Cell 2 for inter-cell separations *d* = 0.1 *µ*m, 1 *µ*m, and 10 *µ*m. The results show monotonic error reduction with compartment refinement, with the induced response being most challenging to resolve at the smallest separations.

The passive-cell results in Fig. 7 display the same overall separation between the directly stimulated and induced responses. The error in Cell 1 remains substantially below that of Cell 2, while the *d* = 0.1 *µ*m and *d* = 1 *µ*m cases exhibit the largest errors. The *d* = 10 *µ*m interaction is considerably better resolved over the discretizations considered. Together, these results confirm that the spatial resolution requirement is governed primarily by the precision needed to capture the field-mediated response of neighboring cells, rather than by the response of the directly stimulated cell.

**Figure 7.**
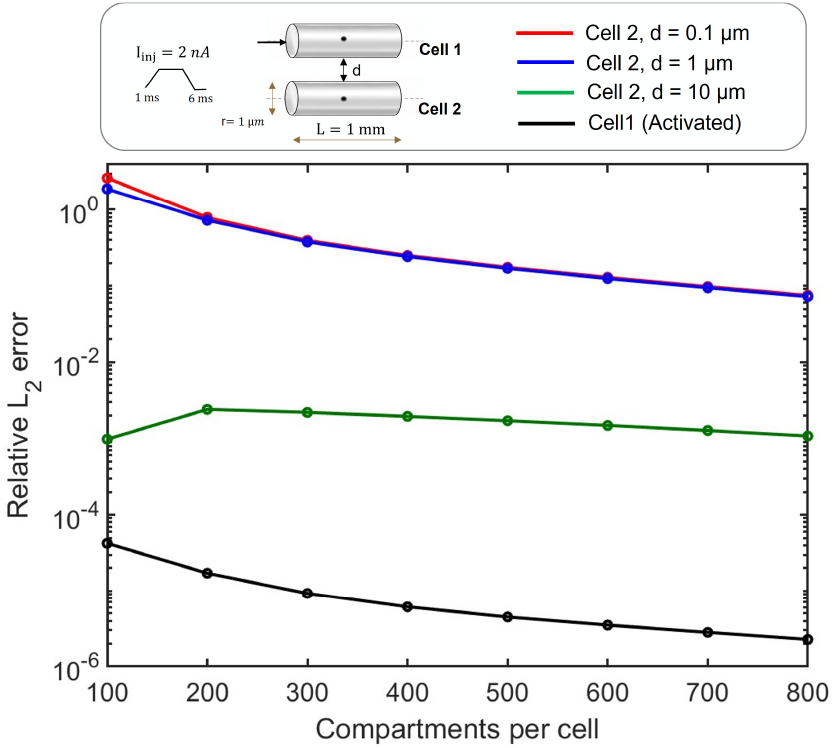
Spatial convergence of the Cable-BEM solution for two parallel passive cylinders. Relative *L*_2_ errors were computed with respect to the reference solution obtained using 1600 compartments per cell. The black curve shows the error in the stimulated cell (Cell 1), while the colored curves show the error in the induced membrane response of Cell 2 for inter-cell separations *d* = 0.1 *µ*m, 1 *µ*m, and 10 *µ*m. As in the active case, the error decreases as the number of compartments per cell increases, and the strongest coupling cases (*d* = 0.1 *µ*m and 1 *µ*m) exhibit the largest errors.

#### 3.2.2 ACA/HODLR Accuracy and Compression

To characterize the low-rank structure of the Cable-BEM operator, a 5 × 5 lattice of 25 parallel cells was considered, featuring 200 compartments per cell and a fixed ACA/HODLR tolerance of 10^*−*7^. The packing density was varied using volume filling fractions (VFFs) of 10.0%, 15.4%, 18.6%, 21.4%, 24.8%, and 31.7%.

Figure 8 illustrates the corresponding HODLR rank distributions. The maximum rank decreases slightly from 136 to 127 as the VFF increases. Despite the reduction in inter-cell spacing, the rank distributions remain highly consistent across all packing densities. The mean rank remains close to 23, while the median rank remains stable at 12. Thus, the hierarchical low-rank structure of the Cable-BEM operator is robustly preserved over the tested range of packing densities.

**Figure 8.**
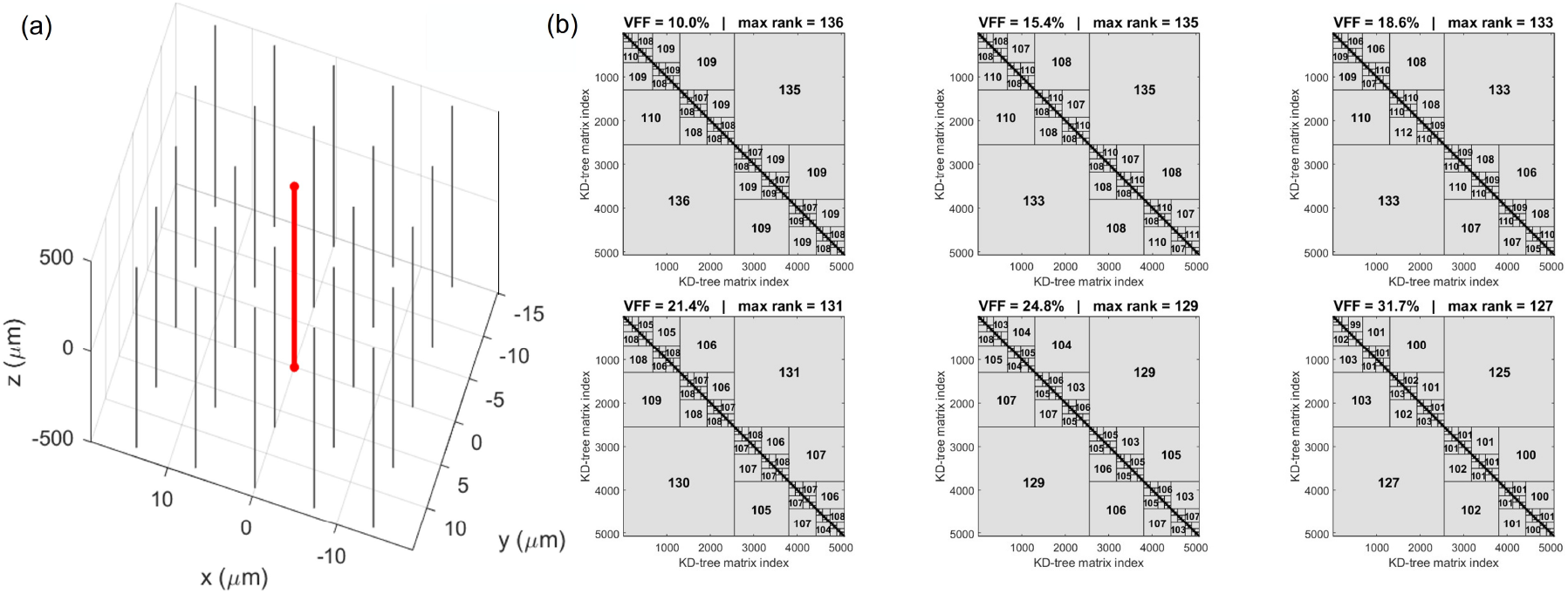
HODLR rank distributions for a 5 *×* 5 lattice of 25 parallel cells with 200 compartments per cell (*N*_red_ = 5075) and a fixed ACA/HODLR tolerance of 10^*−*7^. Panels (a)–(f) correspond to volume filling fractions of 10.0%, 15.4%, 18.6%, 21.4%, 24.8%, and 31.7%, respectively. Numerical labels indicate the ranks of the ACA-compressed off-diagonal blocks at the displayed hierarchy levels.

To quantify the effect of compression on solution accuracy, the ACA/HODLR tolerance was varied from 10^*−*3^ to 10^*−*7^ for a fixed 5 × 5 lattice with 200 compartments per cell (*N*_red_ = 5075) and a volume filling fraction of 18.6%. The resulting compressed solutions were then compared against the dense reference solution.

Figure 9(a) demonstrates that tightening the tolerance systematically improves agreement with the dense reference solution. The maximum relative error decreases from approximately 2.37 × 10^*−*1^ at 10^*−*3^ to below 10^*−*5^ at 10^*−*7^, with the most significant reduction occurring between 10^*−*3^ and 10^*−*4^. The mean relative error follows an identical trend.

**Figure 9.**
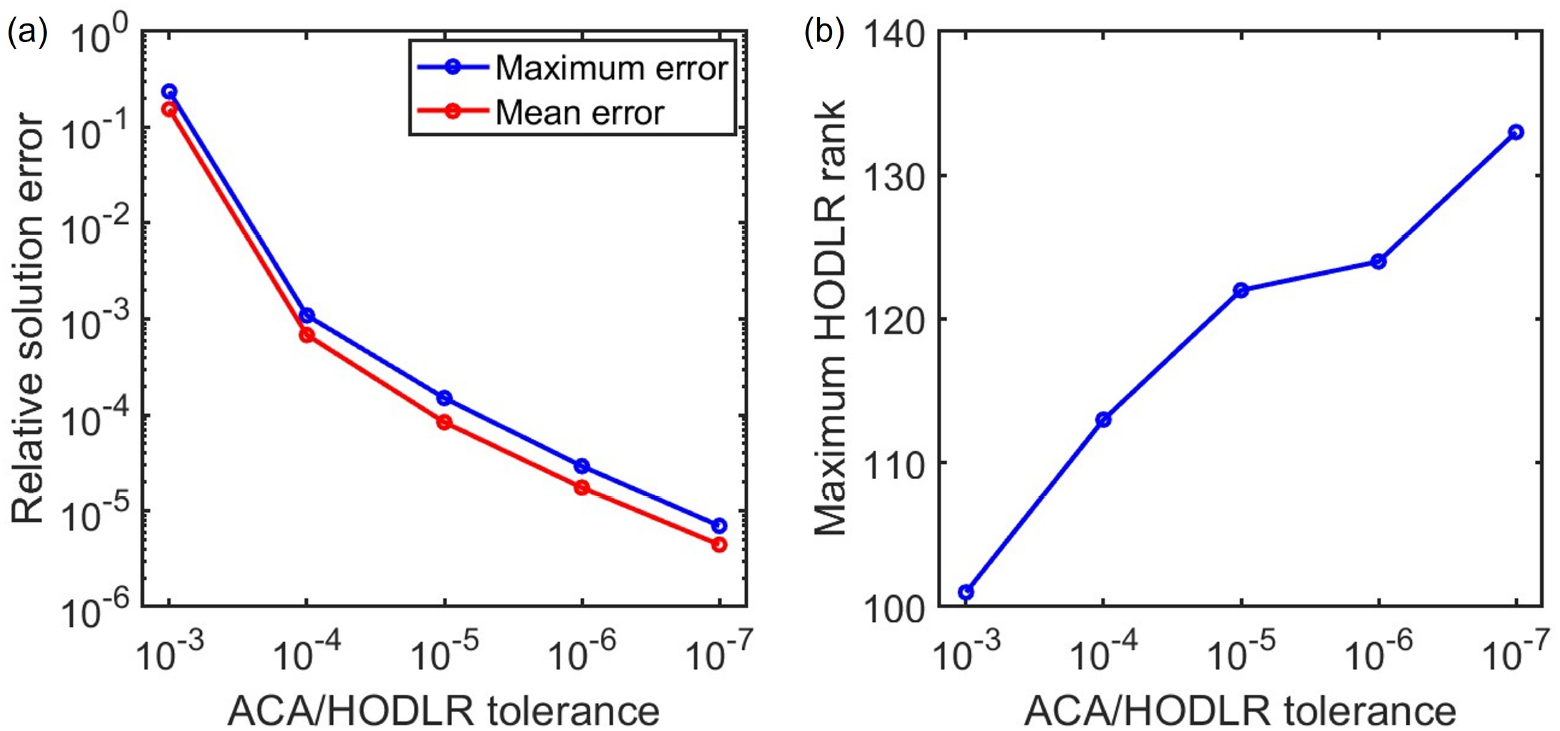
Effect of the ACA/HODLR tolerance on solution accuracy and compression rank for a fixed 5 *×* 5 lattice of 25 cells with 200 compartments per cell (*N*_red_ = 5075) and a volume filling fraction of 18.6%. (a) Maximum and mean relative solution errors with respect to the dense reference solution. (b) Maximum HODLR rank.

This accuracy improvement necessitates a corresponding increase in the rank required for compression, as depicted in Fig. 9(b). The maximum HODLR rank scales from 101 at a tolerance of 10^*−*3^ to 133 at 10^*−*7^. Over this same range, the compressed storage requirement increases from approximately 41 MB to 55 MB. Consequently, tightening the ACA tolerance yields a clear accuracy– compression trade-off: substantially enhanced solution precision is achieved at the expense of a moderate increase in rank and storage me mory.

#### 3.2.3 Effect of Cell Packing

To determine whether denser cell packing compromises the computational efficiency of the HODLR representation, the volume filling fraction (VFF) was varied for a fixed 5 × 5 lattice of 25 cells. Six VFFs were evaluated: 10.0%, 15.4%, 18.6%, 21.4%, 24.8%, and 31.7%, corresponding to neighboring surface-to-surface separations of approximately 4.51, 3.15, 2.64, 2.29, 1.95, and 1.44 *µ*m, respectively. Each cell was discretized with 200 compartments, and the ACA/HODLR tolerance was maintained at 10^*−*7^.

Across these six packing configurations, the compressed storage footprint remains nearly constant, varying narrowly between 52 MB and 55 MB. The HODLR construction, LU factorization, and mean solution times also remain stable within ranges of 851–1261 s, 3.0–7.7 s, and 0.030–0.041 s, respectively, displaying no systematic scaling penalty as the VFF is increased. Therefore, over the investigated spatial range, decreasing the separation between neighboring cells does not meaningfully inflate the storage requirements or computational overhead of the compressed Cable-BEM operator.

#### 3.2.4 Computational Scaling

To evaluate how computational demands scale as the multicellular system expands, the lattice size was progressively increased from 3 ×3 to 15 ×15, corresponding to arrays of 9, 25, 49, 100, and 225 cells. Each cell was discretized with 200 compartments, resulting in reduced system sizes of *N*_red_ = 1827, 5075, 9947, 20300, and 45675, respectively. The volume filling fraction was fixed at 18.6%, and the ACA/HODLR tolerance was held at 10^*−*7^.

Figure 10(a) plots the maximum HODLR rank as the number of cells increases. The maximum rank grows substantially across the evaluated range, reaching 1105 for the 15 × 15 lattice. This trend indicates that increasingly large off-diagonal blocks demand higher numerical ranks as the physical footprint of the multicellular system expands.

**Figure 10.**
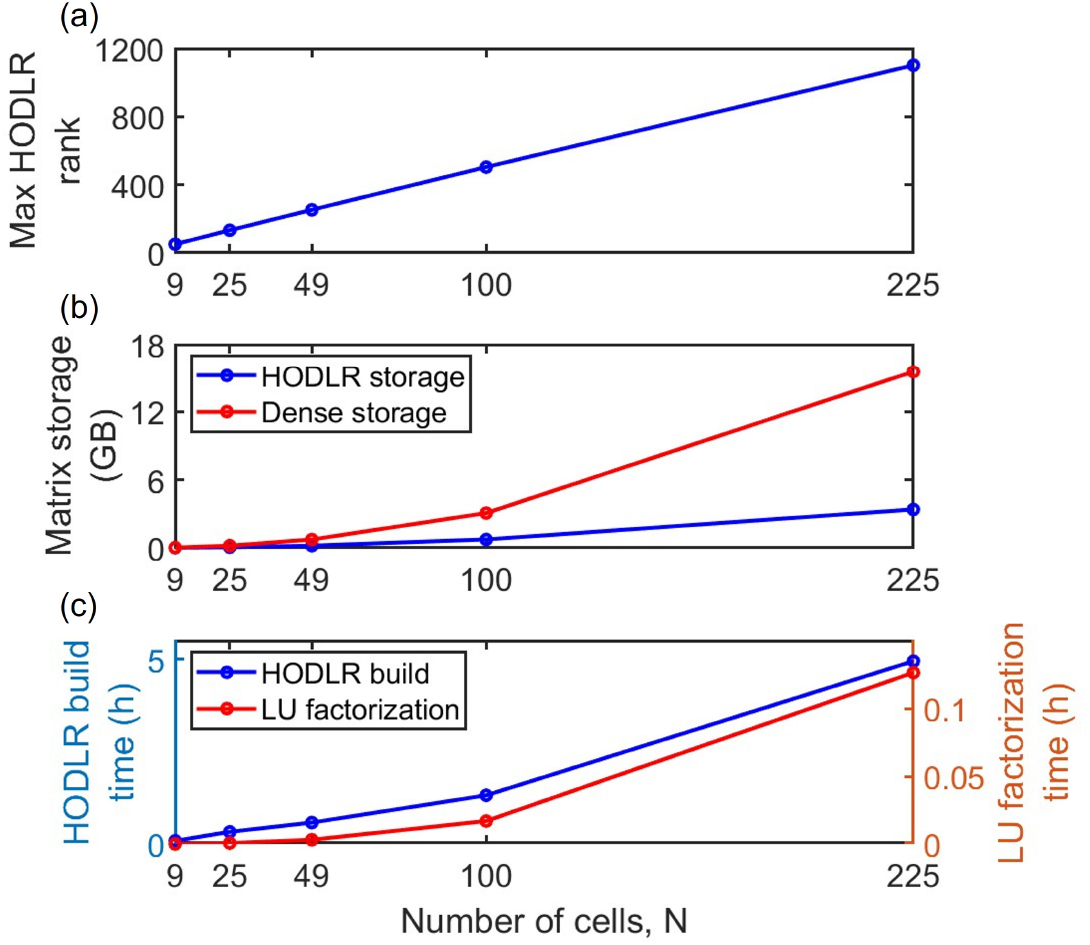
Computational scaling of the ACA-HODLR representation for lattices containing 9, 25, 49, 100, and 225 cells, with 200 compartments per cell, a volume filling fraction of 18.6%, and an ACA/HODLR tolerance of 10^*−*7^. (a) Maximum HODLR rank as a function of the number of cells. (b) Storage required by the compressed HODLR representation compared with the equivalent dense matrix storage. (c) HODLR construction and LU factorization times.

Despite this rank growth, the compressed representation continues to afford substantial memory savings relative to the equivalent dense matrix formalism (Fig. 10(b)). For the largest simulated geometry (225 cells, *N*_red_ = 45675), the HODLR representation requires approximately 3.39 GB of memory, compared to an estimated 15.5 GB for the corresponding dense matrix. This represents a memory footprint reduction by a factor of approximately 4.6.

Figure 10(c) details the HODLR construction and LU factorization times. HODLR construction serves as the dominant computational bottleneck and scales markedly with system size. For the 225-cell case, construction requires approximately 1.84 × 10^4^ s (∼ 5.1 h), whereas the subsequent LU factorization takes only 470 s (∼7.8 min). Crucially, once this factorization is complete, the mean solution time for subsequent steps remains trivial, requiring only 0.38 s for the largest system.

Overall, the current implementation successfully maintains significant matrix compression as the multicellular system is enlarged, though the scaling of the maximum rank and the initial HODLR construction time emerge as the primary computational constraints for massive problem sizes.

To determine whether the observed rank growth is dictated by the total system size or by the geometric nature of the compressed interactions, we further isolated and examined the ACA rank of spatial cluster pairs extracted from a 100 × 100 lattice using the identical geometric partitioning strategy. For each physical box size, two cluster configurations were analyzed: directly adjacent clusters, and clusters separated by one full box width.

As shown in Figure 11, across all three ACA tolerances, the ACA rank of adjacent clusters increases sharply as a function of physical box size, whereas the rank of the separated clusters remains substantially lower and exhibits much weaker spatial scaling. Specifically, for the three physical box sizes of 24.8, 49.6, and 99.1 *µ*m, the adjacent-cluster ranks scale from 49 to 152 and then to 520 at an ACA tolerance of 10^*−*3^, from 79 to 245 and then to 928 at 10^*−*5^, and from 104 to 299 and then to 1071 at 10^*−*7^. Over this same range, the ranks for clusters separated by approximately one box width remain significantly smaller, yielding values of 30, 34, and 33 at 10^*−*3^; 53, 72, and 67 at 10^*−*5^; and 72, 109, and 115 at 10^*−*7^.

**Figure 11.**
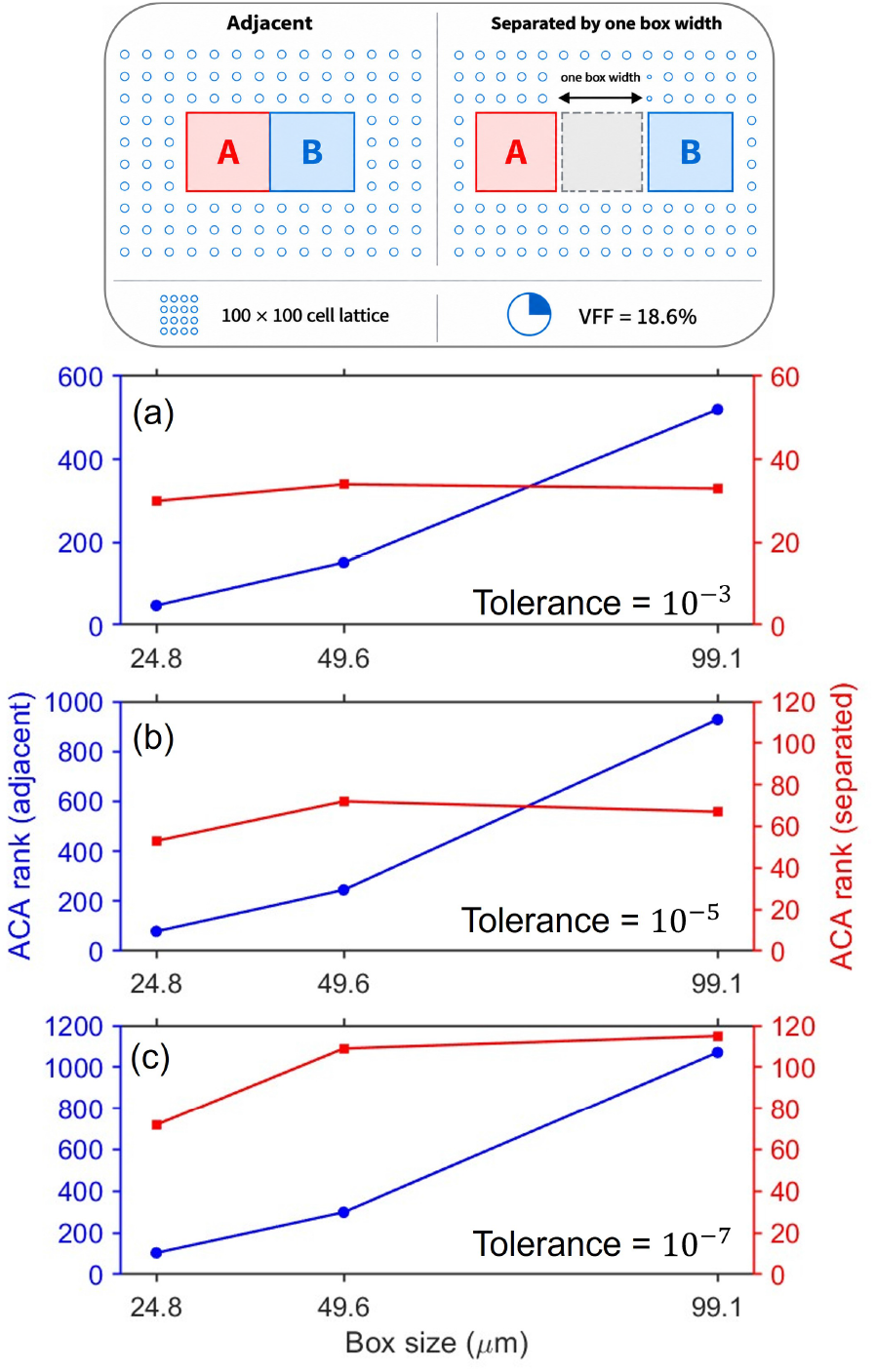
ACA rank as a function of physical box size for adjacent clusters and clusters separated by approximately one box width. The box sizes of 24.8, 49.6, and 99.1 *µ*m contain 180, 1296, and 10944 degrees of freedom, respectively. Panels (a), (b), and (c) correspond to ACA tolerances of 10^*−*3^, 10^*−*5^, and 10^*−*7^, respectively.

The computational overhead and measured ACA errors highlight a similar contrast between the two cluster configurations. For the largest physical box size of 99.1 *µ*m, the adjacent-cluster cases required 1459.7 s, 2758.1 s, and 2731.8 s for tolerances of 10^*−*3^, 10^*−*5^, and 10^*−*7^, respectively, yielding corresponding measured errors of 3.69 × 10^*−*1^, 1.18 × 10^*−*3^, and 3.66 × 10^*−*5^. Conversely, for the separated cluster configuration, the corresponding runtimes were substantially lower—requiring only 419.0 s, 543.3 s, and 971.0 s—while achieving significantly higher precision with measured errors of 4.35 × 10^*−*4^, 4.81 × 10^*−*6^, and 6.39 × 10^*−*8^, respectively. These metrics further confirm that spatially separated interactions are fundamentally more compressible and far less expensive to algebraically approximate.

These findings demonstrate that the rank growth observed in the weak-admissibility HODLR representation is driven primarily by near-field boundary interactions across neighboring cluster interfaces, rather than by the absolute number of unknowns. This further implies that adopting a strong-admissibility hierarchical scheme—where near-neighbor interactions are strictly retained as dense near-field blocks and low-rank compression is applied exclusively to sufficiently separated clusters—could effectively mitigate this specific rank-growth limitation in future solver implementations.

### 3.3 Ephaptic Coupling of Two Cylindrical Cells

Having established the spatial resolution required to accurately resolve the field-mediated response of neighboring cells, we next evaluated the ephaptic interaction itself using a canonical two-cell cylindrical configuration. Two parallel Hodgkin–Huxley cells of length *L* = 1 mm and radius *r* = 1 *µ*m were simulated at surface-to-surface separations of *d* = 0.1, 1, and 10 *µ*m. Cell 1 was stimulated by a 90 pA intracellular current injection, while Cell 2 received no direct active stimulation.

Figure 12(a) traces the membrane potential at the center of the stimulated cell. The core action-potential waveform remains essentially unchanged across the three intercellular separations. By contrast, Fig. 12(b) highlights a transient membrane-potential perturbation induced in the unstimulated Cell 2. This induced response falls on the order of a few microvolts and scales inversely with distance, peaking at the smallest separations, while the *d* = 10 *µ*m geometry exhibits a markedly attenuated perturbation. Consequently, although the extracellular interaction induces only a minor membrane voltage perturbation compared to the source action potential, the Cable BEM formulation successfully resolves a distinct, distance-dependent ephaptic response in the neighboring cell. This simplified geometry effectively isolates the underlying field-mediated physical interaction between two active cells.

**Figure 12.**
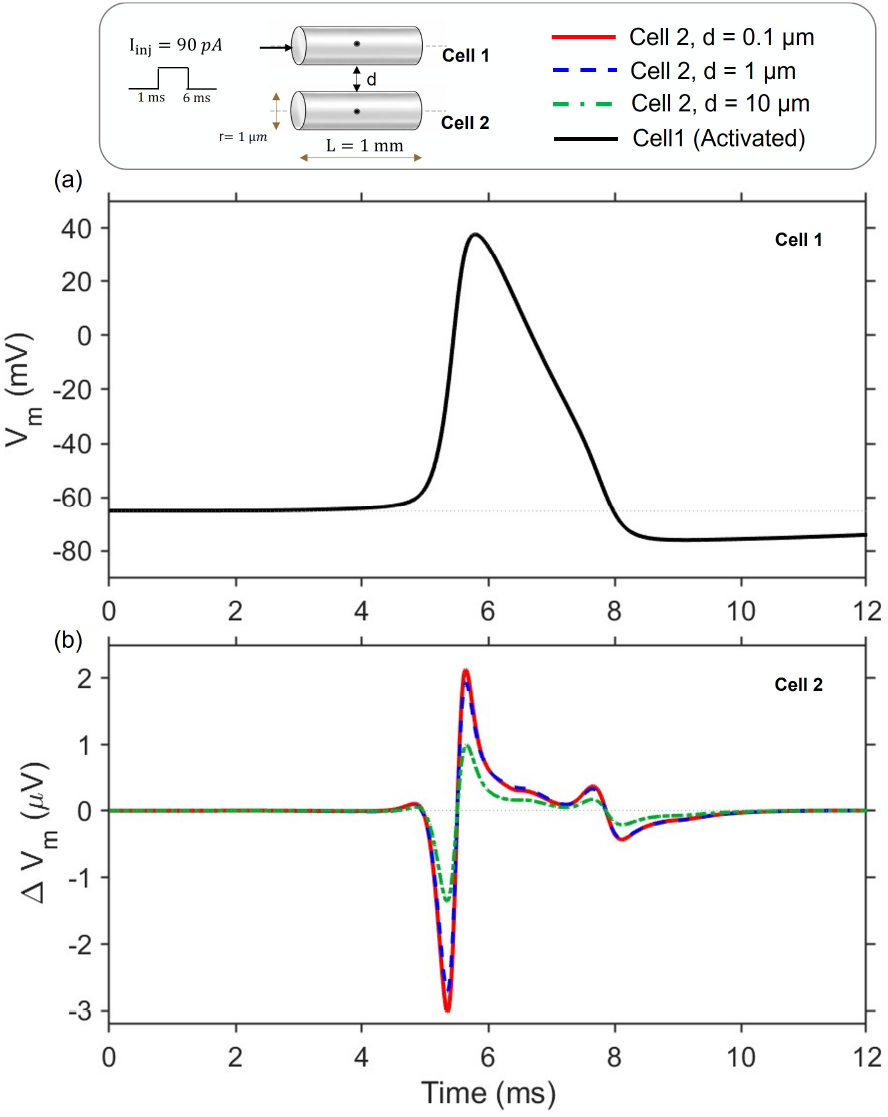
Ephaptic coupling between two parallel Hodgkin–Huxley cylindrical cells. Cell 1 is stimulated by a 90 pA intracellular current injection, while Cell 2 receives no direct stimulation. The cells have length *L* = 1 mm and radius *r* = 1 *µ*m, with intercellular separations *d* = 0.1, 1, and 10 *µ*m. (a) Membrane potential *V*_*m*_ at the center of the stimulated Cell 1. (b) Induced membrane-potential change Δ*V*_*m*_ at the center of the neighboring Cell 2 for the three intercellular separations.

### 3.4 Ephaptic Synchronization of Purkinje Cells

To investigate these ephaptic interactions within a realistic neuronal morphology, we implemented a cerebellar Purkinje cell model based on the detailed morphology, membrane dynamics, and region-specific ion-channel distributions documented by Jæger and Tveito [20]. The corresponding numerical model is publicly available via the Zenodo repository associated with their study. Two neighboring Purkinje cells were simulated across varying inter-axonal separations to quantify the specific influence of ephaptic coupling on their relative firing synchronization.

Figure 13(a) plots the action-potential (AP) lag—defined as the temporal difference between the firing times of Cell 2 and Cell 1—as a function of time. In the uncoupled benchmark Cable model, the AP lag remains approximately constant throughout the simulation, confirming that the initial timing offset between the two cells is artificially preserved. In contrast, the coupled Cable BEM simulations reveal a progressive decrease in the AP lag, directly demonstrating ephaptically mediated phase synchronization. The rate of synchronization exhibits a strong dependence on interaxonal separation: cells separated by just 1 *µ*m approach synchrony most rapidly, while increasing the physical separation progressively attenuates the rate of AP lag decay. At the largest spatial separation evaluated, the field interaction is sufficiently diluted that only a gradual, marginal drift in the AP lag is observable over the simulated temporal window.

**Figure 13.**
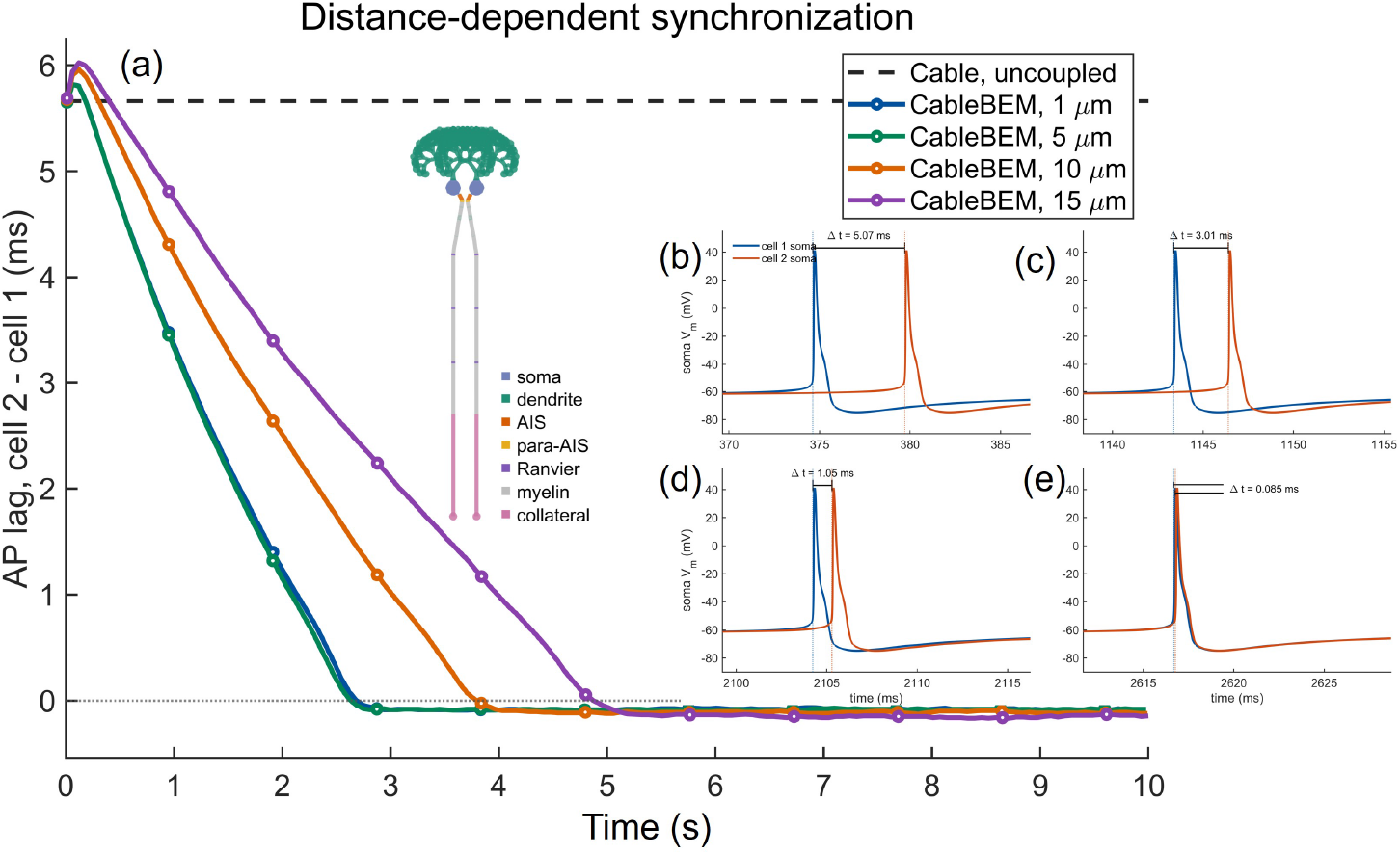
Ephaptic synchronization between two Purkinje cells as a function of inter-axonal separation. **(a)** Action-potential (AP) lag, defined as the difference between the firing times of Cell 2 and Cell 1, for the uncoupled Cable model and the coupled Cable BEM model at separations of 1, 5, 10, and 15 *µ*m. The uncoupled model preserves an approximately constant AP lag, whereas Cable BEM predicts a progressive reduction in lag, with faster synchronization at smaller separations. **(b)–(e)** Representative soma membrane-potential traces for the 1 *µ*m case at successive times, illustrating the progressive reduction in spike-time difference as the two cells approach synchrony.

The temporal evolution of this underlying synchronization process is visualized in Fig. 13(b)–(e) for the 1 *µ*m inter-axonal separation scenario. Initially, the soma membrane potentials are clearly offset in time, reflecting the externally imposed difference in their firing phases. As the cells continue to fire, the temporal discrepancy between corresponding sequential action potentials progressively narrows until the two waveforms become nearly coincident. These dynamics decisively illustrate that the self-consistent extracellular coupling captured by the Cable BEM formulation is capable of driving progressive phase synchronization between biophysically realistic neuronal models, with the coupling strength robustly governed by the intercellular distance.

## 4 Conclusion

We introduced Cable-BEM, a cable-compatible bidomain boundary-element formulation that retains the compartmental degrees of freedom of conventional cable models while solving for the intra- and extracellular fields self-consistently. By analytically incorporating the circumferential dependence of cylindrical membrane elements, the formulation avoids the explicit surface discretization required by full bidomain BEM while preserving nonlocal, field-mediated interactions between neuronal compartments.

Validation against established full bidomain-BEM benchmarks demonstrated robust agreement for intracellular current injection, externally applied electric fields, and multicell field interactions. Spatial-convergence studies further revealed that the field-mediated response induced in a neighboring cell is more computationally demanding to resolve than the response of the directly stimulated cell, particularly at small intercellular separations. These results underscore the importance of rigorously verifying the convergence of key quantities when modeling weak ephaptic effects.

The dense Cable-BEM operator was also examined using ACA/HODLR compression. Our results show that this hierarchical representation can substantially reduce matrix storage while maintaining controllable accuracy across the tested configurations. Concurrently, the observed increase in numerical rank and HODLR construction cost with system size indicates that the computational scaling of the method warrants further investigation and optimization.

Finally, the formulation successfully captured distance-dependent ephaptic interactions between parallel active cylindrical cells and progressive ephaptic synchronization between morphologically realistic Purkinje cells. These results demonstrate that small extracellular perturbations generated by neuronal activity can be resolved self-consistently and can accumulate to produce measurable changes in firing dynamics. Overall, the Cable-BEM method provides a powerful framework for studying field-mediated interactions in multicellular neuronal systems, retaining a computational representation that is closely aligned with conventional cable models.

### A Elliptic-integral treatment of near-field interactions

For nearby cable elements, direct tensor-product quadrature converges slowly because the Green’s-function kernels vary rapidly as the source and observation surfaces approach one another. In the present implementation, element pairs are treated as near-field interactions when they belong to neighboring compartments of the same cell or when their centroid separation is below 10 *µ*m. For these interactions, the source azimuthal coordinate is integrated analytically, while the remaining source-axial and observation-surface integrations are evaluated numerically.

We utilize the cable-element parameterization defined in Eq. (8). At each source coordinate *s*′, the element contains a circular ring of radius *a* = *a*_*e*_(*s*′) centered at 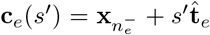. Thus, the source surface integral separates into an axial integral over *s*′ and an azimuthal integral around the corresponding ring. We next evaluate these azimuthal integrals for a fixed observation point **r** and source ring.

For convenience, we introduce a local cylindrical frame centered on the source ring, with 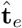 along its axis and 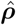 directed toward the radial projection of the observation point. In this frame, the coordinate offsets are written as 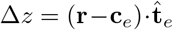 and 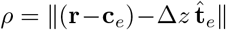, such that the observation point is parameterized as 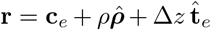. Defining the orthogonal direction 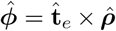, a point on the source ring is represented as 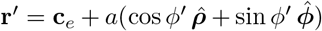. The source–observation distance is therefore given by *R*(*ϕ*′) = [*ρ*^2^ + *a*^2^ −2*aρ* cos *ϕ*′ + Δ*z*^2^]^1*/*2^.

Substituting this parameterization into the source integrals of Eq. (11), the azimuthal integrals required for the adjoint double-layer and hypersingular terms are expressed as

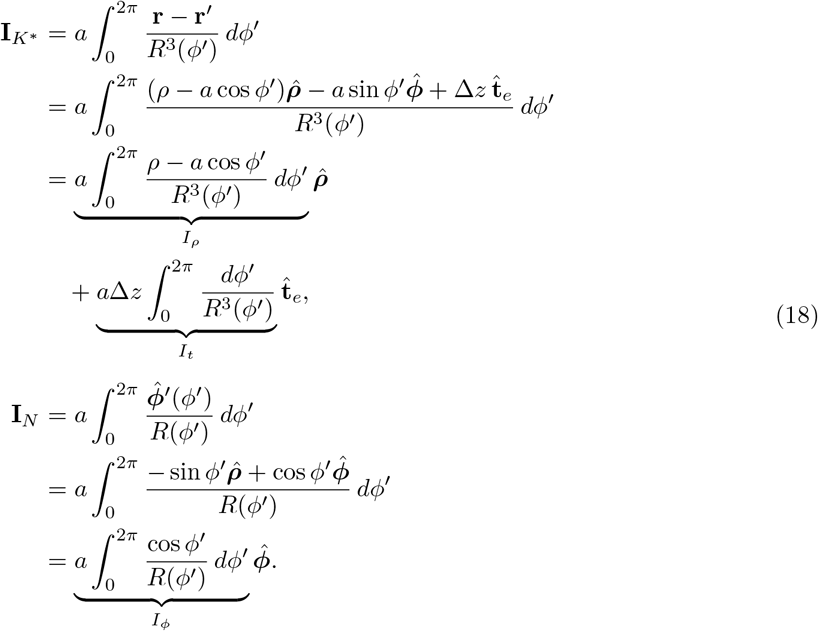

The terms proportional to sin *ϕ*′ vanish by symmetry. To evaluate the remaining integrals, we introduce the half-angle variable *θ* = (*π* − *ϕ*′)*/*2. Thus, we have cos *ϕ*′ = 2 sin^2^ *θ* − 1, and symmetry reduces each azimuthal integral to four times an equivalent integral over 0 ≤ *θ* ≤ *π/*2. Under this transformation, the distance relation is written as 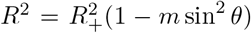, where *R*_*±*_ = [(*a* ± *ρ*)^2^ + Δ*z*^2^]^1*/*2^ denote the maximum and minimum source–observation distances around the ring, respectively, and 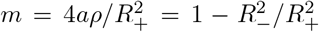 represents the normalized difference between their squared values.

The resulting expressions take the standard form of complete elliptic integrals. Using the definitions 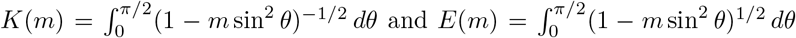, and applying the corresponding identities yields

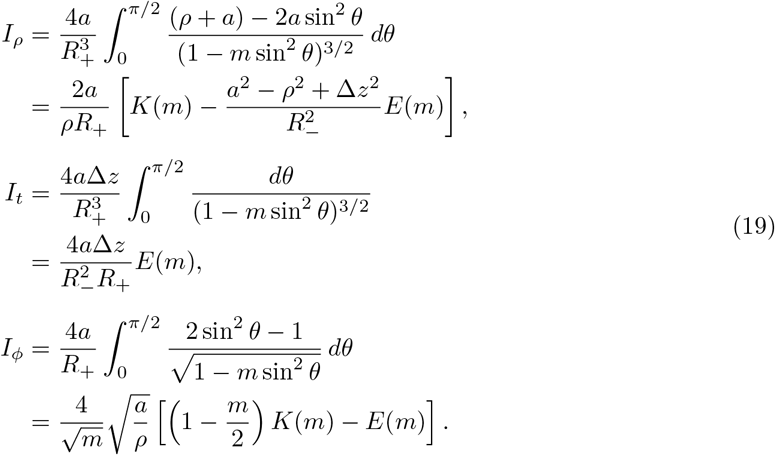

Substituting these analytic ring integrals into the element-interaction formulations completely eliminates the source azimuthal coordinate. For an observation element *e*_*o*_ and source element *e*_*s*_, let 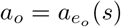 and 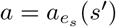. The resulting element-pair contributions are formulated as

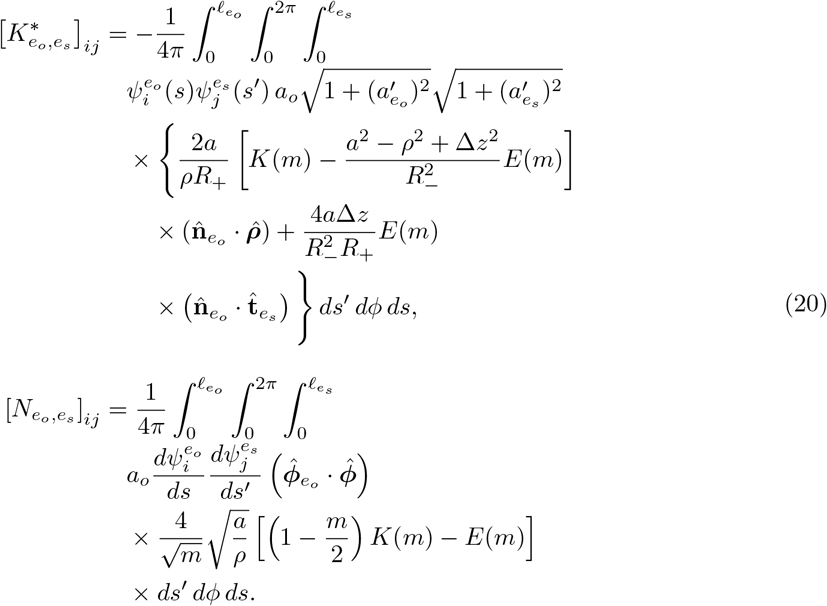

Here, 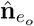 and 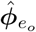 represent the unit normal and azimuthal directions of the observation element, while 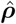 and 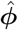 belong to the local cylindrical frame of the source ring. In the adjoint double-layer term, the two square-root factors arise from the observation and source surface metrics; the source metric factor *a dϕ*′ has already been incorporated into *I*_*ρ*_ and *I*_*t*_. In the hypersingular term, the corresponding taper metric factors cancel those arising from 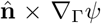, while the factor *a dϕ*′ is already included in *I*_*ϕ*_.

Consequently, this analytical treatment eliminates the source azimuthal integration, leaving the source-axial coordinate *s*′ and the observation coordinates (*s, ϕ*) to be evaluated numerically. For near-field interactions, the observation element is evaluated using 64 axial and 64 azimuthal quadrature points, while the source-axial integral is computed using a 40-point Gaussian quadrature scheme. In the limit *ρ* → 0, we employ the expressions *I*_*ρ*_ = *I*_*ϕ*_ = 0 and *I*_*t*_ = 2*πa*Δ*z/*(*a*^2^ + Δ*z*^2^)^3*/*2^. Mathematically, values of *m* approaching unity are bounded strictly below one to prevent evaluation exactly at the logarithmic singularity of *K*(*m*) at *m* = 1. Well-separated element pairs are instead computed using the conventional, cached tensor-product quadrature employed elsewhere in the solver. This hybrid scheme successfully resolves nearly singular near-field interactions while preserving the underlying cable-element basis and the unified BEM matrix structure of the formulation.

## Acknowledgments

The authors gratefully acknowledge Aman Aberra, Weng Cho Chew, and Yang Liu for their insightful discussions and valuable feedback. This research was sponsored by the U.S. Army Research Laboratory (ARL) and was accomplished under grant numbers W911NF261A103 and W911NF261A083. The views and conclusions contained in this document are those of the authors and should not be interpreted as representing the official policies, either expressed or implied, of the Army Research Laboratory or the U.S. Government.

## Author Contributions

**Vanine Sabino:** Conceptualization, methodology, software development, writing (original draft preparation), data curation, formal analysis, and visualization. **Amanda Walenciak:** Neuron models, visualization, and editing. All authors discussed the results and contributed to the final manuscript. **Luis J. Gomez:** Conceptualization, methodology, software development, visualization, writing, supervision, and funding acquisition.

## Declarations

### Ethical Statement

The authors declare that this study does not involve any human participants or animal experiments.

### Conflict of Interest

The authors declare that they have no known competing financial interests or personal relationships that could have appeared to influence the work reported in this paper.

### Funding

This research was sponsored by the U.S. Army Research Laboratory (ARL) and was accomplished under grant numbers W911NF261A103 and W911NF261A083. The views and conclusions contained in this document are those of the authors and should not be interpreted as representing the official policies, either expressed or implied, of the Army Research Laboratory or the U.S. Government.

### Data Availability Statement

The data that support the findings of this study are available from the corresponding author upon reasonable request.

## Notes

### Competing Interest Statement

The authors have declared no competing interest.

## References

[1] V. Sabino, A. Murugesan, N. I. Hasan, S. S. Vaezi, A. Walenciak, P. Jayatissa, and L. J. Gomez, “Advances in computational electromagnetics for enhanced noninvasive brain stimulation: E-field dosimetry, uncertainty quantification, optimization, and neural response modeling,” IEEE Antennas and Propagation Magazine, 2026.

[2] A. V. Peterchev, T. A. Wagner, P. C. Miranda, M. A. Nitsche, W. Paulus, S. H. Lisanby, A. Pascual-Leone, and M. Bikson, “Fundamentals of transcranial electric and magnetic stimulation dose: definition, selection, and reporting practices,” Brain stimulation, vol. 5, no. 4, pp. 435–453, 2012.

[3] M. Bikson, A. Rahman, and A. Datta, “Computational models of transcranial direct current stimulation,” Clinical EEG and neuroscience, vol. 43, no. 3, pp. 176–183, 2012.

[4] C. C. McIntyre, W. M. Grill, D. L. Sherman, and N. V. Thakor, “Cellular effects of deep brain stimulation: model-based analysis of activation and inhibition,” Journal of neurophysiology, 2004.

[5] C. A. Anastassiou and C. Koch, “Ephaptic coupling to endogenous electric field activity: why bother?” Current opinion in neurobiology, vol. 31, pp. 95–103, 2015.

[6] C. A. Anastassiou, S. M. Montgomery, M. Barahona, G. Buzsáki, and C. Koch, “The effect of spatially inhomogeneous extracellular electric fields on neurons,” Journal of Neuroscience, vol. 30, no. 5, pp. 1925–1936, 2010.

[7] J. H. Goldwyn and J. Rinzel, “Neuronal coupling by endogenous electric fields: cable theory and applications to coincidence detector neurons in the auditory brain stem,” Journal of neurophysiology, vol. 115, no. 4, pp. 2033–2051, 2016.

[8] F. Fröhlich and D. A. McCormick, “Endogenous electric fields may guide neocortical network activity,” Neuron, vol. 67, no. 1, pp. 129–143, 2010.

[9] J. Jefferys, “Nonsynaptic modulation of neuronal activity in the brain: electric currents and extracellular ions,” Physiological reviews, vol. 75, no. 4, pp. 689–723, 1995.

[10] A. L. Hodgkin and A. F. Huxley, “A quantitative description of membrane current and its application to conduction and excitation in nerve,” The Journal of Physiology, vol. 117, no. 4, pp. 500–544, 1952.

[11] M. L. Hines and N. T. Carnevale, “The NEURON simulation environment,” Neural Computation, vol. 9, no. 6, pp. 1179–1209, 1997.

[12] F. Rattay, “Analysis of models for external stimulation of axons,” IEEE Transactions on Biomedical Engineering, vol. BME-33, no. 10, pp. 974–977, 1986.

[13] D. R. McNeal, “Analysis of a model for excitation of myelinated nerve,” IEEE Transactions on Biomedical Engineering, vol. BME-23, no. 4, pp. 329–337, 1976.

[14] C. Gold, D. A. Henze, C. Koch, and G. Buzsáki, “On the origin of the extracellular action potential waveform: A modeling study,” Journal of Neurophysiology, vol. 95, no. 5, pp. 3113–3128, 2006.

[15] G. R. Holt and C. Koch, “Electrical interactions via the extracellular potential near cell bodies,” Journal of Computational Neuroscience, vol. 6, no. 2, pp. 169–184, 1999.

[16] A. Tveito, K. H. Jæger, G. T. Lines, Ł. Paszkowski, J. Sundnes, A. G. Edwards, T. Māki-Marttunen, G. Halnes, and G. T. Einevoll, “An evaluation of the accuracy of classical models for computing the membrane potential and extracellular potential for neurons,” Frontiers in computational neuroscience, vol. 11, p. 27, 2017.

[17] B. Wang, A. S. Aberra, W. M. Grill, and A. V. Peterchev, “Modified cable equation incorporating transverse polarization of neuronal membranes for accurate coupling of electric fields,” Journal of Neural Engineering, vol. 15, no. 2, p. 026003, 2018.

[18] A. Agudelo-Toro and A. Neef, “Computationally efficient simulation of electrical activity at cell membranes interacting with self-generated and externally imposed electric fields,” Journal of neural engineering, vol. 10, no. 2, p. 026019, 2013.

[19] C. A. Anastassiou, R. Perin, H. Markram, and C. Koch, “Ephaptic coupling of cortical neurons,” Nature Neuroscience, vol. 14, no. 2, pp. 217–223, 2011.

[20] K. H. Jæger and A. Tveito, “Extracellular stimulation and ephaptic coupling of neurons in a fully coupled finite element-based extracellular–membrane–intracellular (EMI) model,” Frontiers in Computational Neuroscience, vol. 20, p. 1755548, 2026.

[21] A. Fellner, A. Heshmat, P. Werginz, and F. Rattay, “A finite element method framework to model extracellular neural stimulation,” Journal of Neural Engineering, vol. 19, no. 2, p. 022001, 2022.

[22] S. Joucla, A. Glière, and B. Yvert, “Current approaches to model extracellular electrical neural microstimulation,” Frontiers in computational neuroscience, vol. 8, p. 13, 2014.

[23] W. Ying and C. S. Henriquez, “Hybrid finite element method for describing the electrical response of biological cells to applied fields,” IEEE transactions on biomedical engineering, vol. 54, no. 4, pp. 611–620, 2007.

[24] K. H. Jæger, K. G. Hustad, X. Cai, and A. Tveito, “Efficient numerical solution of the emi model representing the extracellular space (e), cell membrane (m) and intracellular space (i) of a collection of cardiac cells,” Frontiers in Physics, vol. 8, p. 579461, 2021.

[25] H. Meffin, B. Tahayori, D. B. Grayden, and A. N. Burkitt, “Modeling extracellular electrical stimulation: I. derivation and interpretation of neurite equations,” Journal of neural engineering, vol. 9, no. 6, p. 065005, 2012.

[26] B. Tahayori, H. Meffin, S. Dokos, A. N. Burkitt, and D. B. Grayden, “Modeling extracellular electrical stimulation: Ii. computational validation and numerical results,” Journal of neural engineering, vol. 9, no. 6, p. 065006, 2012.

[27] M. S. Hamalainen and J. Sarvas, “Realistic conductivity geometry model of the human head for interpretation of neuromagnetic data,” IEEE transactions on biomedical engineering, vol. 36, no. 2, pp. 165–171, 1989.

[28] S. N. Makarov, L. Golestanirad, W. A. Wartman, B. T. Nguyen, G. M. Noetscher, J. P. Ahveninen, K. Fujimoto, K. Weise, and A. R. Nummenmaa, “Boundary element fast multipole method for modeling electrical brain stimulation with voltage and current electrodes,” Journal of neural engineering, vol. 18, no. 4, p. 0460d4, 2021.

[29] S. N. Makarov, G. M. Noetscher, T. Raij, and A. Nummenmaa, “A quasi-static boundary element approach with fast multipole acceleration for high-resolution bioelectromagnetic models,” IEEE transactions on biomedical engineering, vol. 65, no. 12, pp. 2675–2683, 2018.

[30] D. M. Czerwonky, A. S. Aberra, and L. J. Gomez, “A boundary element method of bidomain modeling for predicting cellular responses to electromagnetic fields,” Journal of neural engineering, vol. 21, no. 3, p. 036050, 2024.

[31] N. I. Hasan, V. Sabino, A. J. Walenciak, Y. Liu, and L. J. Gomez, “Modeling pyramidal neurons using bidomain bem and hierarchical matrix approximation,” bioRxiv, pp. 2025–02, 2025.

[32] W. Hackbusch, “A sparse matrix arithmetic based on-matrices. part i: Introduction to-matrices,” Computing, vol. 62, no. 2, pp. 89–108, 1999.

[33] W. Hackbusch et al., Hierarchical matrices: algorithms and analysis. Springer Heidelberg, 2015, vol. 49.

[34] M. Bebendorf and S. Rjasanow, “Adaptive low-rank approximation of collocation matrices,” Computing, vol. 70, no. 1, pp. 1–24, 2003.

[35] S. Rjasanow, “Adaptive cross approximation of dense matrices,” in Int. Association Boundary Element Methods Conf., IABEM, 2002, pp. 28–30.

[36] S. Kurz, O. Rain, and S. Rjasanow, “The adaptive cross-approximation technique for the 3d boundary-element method,” IEEE transactions on Magnetics, vol. 38, no. 2, pp. 421–424, 2002.

[37] K. Zhao, M. N. Vouvakis, and J.-F. Lee, “The adaptive cross approximation algorithm for accelerated method of moments computations of emc problems,” IEEE transactions on electro-magnetic compatibility, vol. 47, no. 4, pp. 763–773, 2005.

[38] Y. Liu, W. Sid-Lakhdar, E. Rebrova, P. Ghysels, and X. S. Li, “A parallel hierarchical blocked adaptive cross approximation algorithm,” The International Journal of High Performance Computing Applications, vol. 34, no. 4, pp. 394–408, 2020.

[39] D. Wang, N. I. Hasan, M. Dannhauer, A. C. Yucel, and L. J. Gomez, “Fast computational e-field dosimetry for transcranial magnetic stimulation using adaptive cross approximation and auxiliary dipole method (aca-adm),” NeuroImage, vol. 267, p. 119850, 2023.

[40] W. Rall, “Core conductor theory and cable properties of neurons,” Comprehensive Physiology, pp. 39–97, 1977.

[41] W. Rall, “Branching dendritic trees and motoneuron membrane resistivity,” Experimental neurology, vol. 1, no. 5, pp. 491–527, 1959.

[42] W. Rall, “Theory of physiological properties of dendrites,” Annals of the New York Academy of Sciences, vol. 96, no. 4, pp. 1071–1092, 1962.

[43] V. Sabino and L. J. Gomez, “A matlab-based solver for modeling neurons,” in 2025 International Applied Computational Electromagnetics Society Symposium (ACES). IEEE, 2025, pp. 1–2.

[44] N. T. Carnevale and M. L. Hines, The NEURON book. Cambridge University Press, 2006.

[45] F. Rattay, “The basic mechanism for the electrical stimulation of the nervous system,” Neuroscience, vol. 89, no. 2, pp. 335–346, 1999.

[46] D. R. Wilton and N. J. Champagne, “Evaluation and integration of the thin wire kernel,” IEEE transactions on antennas and propagation, vol. 54, no. 4, pp. 1200–1206, 2006.

